# Genetic diversity of the *LILRB1* and *LILRB2* coding regions in an admixed Brazilian population sample

**DOI:** 10.1101/2021.04.16.440206

**Authors:** Maria Luiza Guimarães de Oliveira, Erick C. Castelli, Luciana C. Veiga-Castelli, Alison E. Pereira, Letícia Marcorin, Thássia M. T. Carratto, Andreia S. Souza, Heloisa S. Andrade, Aguinaldo L. Simões, Eduardo A. Donadi, David Courtin, Audrey Sabbagh, Silvana Giuliatti, Celso Teixeira Mendes-Junior

## Abstract

Leukocyte Immunoglobulin (Ig)-like Receptors (LILR) LILRB1 and LILRB2 play a pivotal role in maintaining self-tolerance and modulating the immune response through interaction with classical and non-classical Human Leukocyte Antigen (HLA) molecules. Although both diversity and natural selection patterns over HLA genes have been extensively evaluated, little information is available concerning the genetic diversity and selection signatures on the *LIRB1/2* regions. Therefore, we identified the *LILRB1/2* genetic diversity using next-generation sequencing in a population sample comprising 528 healthy control individuals from São Paulo State, Brazil. We identified 58 *LILRB1* Single Nucleotide Variants (SNVs), which gave rise to 13 haplotypes with at least 1% of frequency. For *LILRB2*, we identified 41 SNVs arranged into 11 haplotypes with frequencies above 1%. We found evidence of either positive or purifying selection on *LILRB1/2* coding regions. Some residues in both proteins showed to be under the effect of positive selection, suggesting that amino acid replacements in these proteins resulted in beneficial functional changes. Finally, we have shown that allelic variation (six and five amino acid exchanges in LILRB1 and LILRB2, respectively) affects the structure and/or stability of both molecules. Nonetheless, LILRB2 has shown higher average stability, with no D1/D2 residue affecting protein structure. Taken together, our findings demonstrate that *LILRB1* and *LILRB2* are highly polymorphic and provide strong evidence supporting the directional selection regime hypothesis.

## Introduction

Leukocyte immunoglobulin-like receptor–1 (LIR-1/CD85j/ILT2/LILRB1) and Leukocyte immunoglobulin-like receptor–2 (LIR-2/CD85d/ILT4/LILRB2) are members of the leukocyte immunoglobulin (Ig)-like receptor (*LIR*) family. *LIR* comprises a family of 13 inhibitory and stimulatory receptors encoded within the leukocyte receptor complex (*LCR*), located at the chromosomal region 19q.13.4, which also includes killer cell inhibitory receptors (*KIRs*), Leukocyte associated Ig-like receptor (*LAIR*) families, and the *Fc*α*R* gene (Brown et al. 2004). Among the 13 members from the LIR family, 11 genes are functional and encode five activating (*LILRA1, 2, 4–6*), five inhibitory (*LILRB1–5*), and one soluble (*LILRA3*) protein forms (Hirayasu and Arase 2015). Structurally, however, they are categorized into group 1 (LILRB1, LILRB2, and LILRA1–3) and group 2 (LILRB3–5 and LILRA4–6), based on conserved residues shared with LILRB1 (Brown et al. 2004).

Alternative mRNA processing gives rise, respectively, to thirteen and eight different protein isoforms from LILRB1 and LILRB2, showing different cell expression profiles. Whereas LILRB1 is widely expressed in monocytes, dendritic cells, B cells, and subsets of Natural Killer (NK) and T cells, LILRB2 expression is restricted within the myelomonocytic lineage (Colonna et al. 1997; Young et al. 2001; Brown et al. 2004). It has been detected the LILRB1/2 expression in osteoclasts and placental stroma, the LILRB1 expression in breast carcinoma, tissue-like memory B cells, and the LILRB2 expression in placental, vascular smooth muscle and human hematopoietic stem cells (Hudson and Allen 2016).

LILRB1 and LILRB2 proteins contain four (Ig) domains (D1-D4), a transmembrane domain, and 3-4 immunoreceptor tyrosine-based inhibitory motifs (ITIMs) in their cytoplasmic tails that interact with tyrosine phosphatases, inhibiting activating signals (Martin et al. 2002; Kang et al. 2016). LILRB1 and LILRB2 bind to a variety of classical and non-classical HLA molecules through two N-terminal extracellular domains (D1 and D2), being this interaction mediated via α3 and β2m domains in the HLA molecule (Colonna et al. 1997; Borges and Cosman 2000; Willcox et al. 2003; Shiroishi et al. 2006). Moreover, LILRB1/2 modulate the CD8+ T cell activation by competition with CD8 for HLA class I binding (Shiroishi et al. 2003). Conversely, D3 and D4 do not take part of the interaction with HLA molecules, but they seem to act as a scaffold for the D1-D2 binding (Wang et al. 2019).

Both receptors interact with different class-I HLAs Ia (HLA-A, -B, and -C) and Ib (HLA-E -F, and -G), although minimal or no binding to HLA-C has been detected (Colonna et al. 1998; Fanger et al. 1998). LILRB1 also binds to the human cytomegalovirus (HCMV) HLA class I homolog (UL18, Cosman et al. 1997), dengue virus (Chan et al. 2014), the damage-associated molecular pattern proteins S100A8/A9 (Wang et al. 2018) and RIFIN proteins expressed by *P. falciparum* (Saito et al. 2018; Harrison et al. 2020). LILRB1 class-I HLA interaction encompasses an interface where 70% of the LILRB1 D1-D2 surface makes contact with the HLA β2M domain (site 1), and an interface involving the recognition of HLA α3 domain by the LILRB1 D1 domain (site 2, Willcox et al. 2003). Although site 1 adopts a highly conserved orientation in HLA-A2-LILRB1 (Willcox et al. 2003), HLA-F-LILRB1 (Dulberger et al. 2017), HLA-G-LILRB2 (Shiroishi et al. 2006), and also UL18-LILRB1 complex structures (Yang and Bjorkman 2008), β2M-HLA-F shows additional contacts, reminiscent of high-affinity recognition UL18-LILRB1 (around 1000-folds higher than that for regular HLAs, Cosman et al. 1997) that might explain why β2M-HLA-F is the highest-affinity ligand for LILRB1 among class-I HLAs molecules (Dulberger et al. 2017). LILRB1–HLA-A2 structure as making contacts with LILRB1 (α3 residues 193–196, 198, and 248) are usually observed in aforementioned interactions, with residues 195-197 conserved in HLA-A, HLA-B, HLA-C, HLA-E, and HLA-F. The most divergent sequence corresponds to HLA-G, which contains amino acid divergences at positions 195 (Ser to Phe) and 197 (His to Tyr, Willcox et al. 2003). These non-conservative changes were proposed as affecting the binding affinity of LILRB1/2 with HLA-G (3-to 4-fold higher) relative to other class-I HLAs molecules (Shiroishi et al. 2003).

Likewise, LILRB2 seems to play important roles in diverse biological activities once it interacts with different ligands besides HLA class I molecules, such as CD1d, angiopoietin-like proteins (Angptls), myelin inhibitors, and β-amyloid (Kang et al. 2016). LILRB2 binding induces the inhibition of FcR-mediated signaling in monocytes, the serotonin release in LILRB2-transfected basophilic cell lines, the axonal suppression regeneration in myelin cells, the interaction with β-amyloid contributing to the Alzheimer’s disease development, as well as the proliferation of the antigen-presenting cells (APCs) as response to Salmonella infection (Kang et al. 2016). Unlike LILRB1, the LILRB2 receptor does not require the presence of β2m when binding to HLA molecules (Shiroishi et al. 2006). Furthermore, LILRB2 binds HLA-G with higher affinity than LILRB1, which was attributed to residues in the binding region on D1D2 LILRB1 (for instance Y76, D80, and R84) that are not conserved in LILRB2 (Q76, R80, and R84) (Shiroishi et al. 2003) and to the different recognition specificities, whereas residues T36 and A38 in LILRB2 recognize 195–197 loop, T38 and T76 in LILRB1 bind only Phe195 in the HLA-G molecule (Shiroishi et al. 2006).

Unlike most of other MHCs, HLA-G has two free cysteine residues (Cys42 and Cys147), which enables the formation of dimers within the α1 domain (Cys42-Cys42) that binds the receptors with a stronger avidity than HLA-G monomers (Shiroishi et al. 2006). HLA-G produces at least seven distinct isoforms by alternative splicing, with four membrane-bound (HLA-G1-G4) and three soluble (HLA-G5-G7) isoforms (Ishitani and Geraghty 1992; Paul et al. 2000), besides four novel isoforms predicted by Tronik-Le Roux et al. (2017). HLA-G1 and -G5 corresponds to the most extensively studied isoforms and present the typical structure found in HLA class I molecules (α1–α2–α3 associated to β2m), being recognized by both LILRB1 and LILRB2 receptors. In contrast, HLA-G2 and -G6 isoforms (α1–α3 domains), bind only to LILRB2 (Furukawa et al. 2019). Except for HLA-G3 and -G7, all isoforms can dimerize. The dimers most frequently detected correspond to HLA-G1 and - G5 dimers (HoWangYin et al. 2012), however, HLA-G2 and -G6 dimers have a functional significance given the existence of individuals with HLA-G–null alleles (HLA-G*0105N) that cannot encode functional HLA-G1 molecules (Menier et al. 2000), which has been shown to bind to LILRB2 with a significant avidity effect (Kuroki et al. 2017).

Compared to the high allelic diversity found in the KIR complex, LILRB1 and LILRB2 are considered moderately polymorphic (Young et al. 2001). Notwithstanding, as other members of the LIR family, they are rapidly evolving and showing larger interspecies differences compared to the genome average (Canavez et al. 2001). Therefore, functional polymorphisms in these genes may influence the immune response diversity among individuals, and consequently, their susceptibility to diseases (Kuroki et al. 2005).

Admixed populations represent a great opportunity to identify new polymorphisms and genetic factors involved in disease susceptibility (Shriner et al. 2011). The Brazilian population is highly admixed, the result of centuries of admixture among three main parental groups: Native Americans, Europeans, and Africans (Salzano and Bortolini 2002). While this intensive admixture brings a methodological issue for genetic analysis, the Brazilian population’s complexity has advantages for disease gene mapping and genetic diversity survey. Among the features that make the Brazilian population particularly valuable for disease *loci* mapping is the higher genetic diversity in comparison with most parental populations and population-specific patterns of linkage disequilibrium, which may modify the length of haplotypes and distribution of haplotype frequencies in the admixed population in comparison with that in the parental populations (Shriner et al. 2011). Notably, the linkage disequilibrium, occasionally maintained over large genetic distances, may also result from natural selection action (Pritchard and Przeworski 2001).

Protein evolution is the outcome of the interplay between the mutational process and selective forces (Montoya-Burgos 2011). Nevertheless, the relationship between sequence and function has proven difficult to be understood. Therefore, we have explored natural selection signatures as proxies for potential changes in LILRB1/2 functions, further assessed by structural protein analysis. Although the genetic diversity and evolutionary dynamics of *HLA* genes have been extensively evaluated, with patterns of balancing (Hughes and Nei 1988; Takahata et al. 1992; Meyer and Thomson 2001; Garrigan and Hedrick 2003) and positive (Sabeti et al. 2006; Bakker et al. 2009) selection well established, few studies have addressed the population genetics aspects of *LILRB1/2* (Hirayasu et al. 2008). Accordingly, this study proposes to describe the genetic diversity pattern of these genes in an admixed Brazilian population sample, besides investigating the evolutionary and functional significance of *LILRB1/2* variants.

## Materials and Methods

### Population sample

A total of 528 unrelated individuals, 266 females and 262 males, aged 18 to 80 years, were enrolled from the blood bank of Ribeirão Preto (Hemocentro de Ribeirão Preto) and University of São Paulo Hospital in Ribeirão Preto (Hospital das Clínicas de Ribeirão Preto), São Paulo state, Southeastern Brazil. All participants signed informed consent forms, following the guidelines provided by the Research Ethics Committee of the Faculdade de Filosofia, Ciências e Letras de Ribeirão Preto, FFCLRP-USP, as approved by the CEP-FFCLRP CAAE n.25696413.7.0000.5407 protocol.

### Laboratory Analysis

Genomic DNA was extracted by standard salting-out protocol (Miller et al. 1988) and stored at -20°C until library preparation. DNA quality was evaluated in terms of purity level, integrity, and concentration by NanoDrop® ND-1000 (Thermo Fisher Scientific Inc., Waltham, MA), Agarose Gel Electrophoresis, and Qubit™ dsDNA BR Assay (Life Technologies), respectively. Finally, samples were normalized to 5ng/µL to achieve a standard concentration for sequencing-library preparation.

SureDesign Tool (Agilent Technologies, Inc., Santa Clara, CA) was used to the design probes to target a total of 284,367bp regions, including the coding DNA sequences (CDS) of *LILRB1* (NM_006669.6) and *LILRB2* (NM_001080978.4). Libraries were prepared using a customized HaloPlex Target Enrichment System (Agilent Technologies, Inc., Santa Clara, CA), following the manufacturer’s instructions. DNA libraries were quantified before sequencing by Qubit® 2.0 Fluorometer (Thermo Fisher Scientific Inc., Waltham, MA) and 2100 Bioanalyzer (Agilent Technologies, Inc., Santa Clara, CA). Lastly, a pool of up to 96 samples was diluted to 4 nmol/L and inserted as input for sequencing using MiSeq Reagent kit V3 (600 cycles), in the MiSeq Personal Sequencer (Illumina Inc., San Diego, CA).

### Bioinformatics Approaches

Sequencing reads were trimmed for adapters sequences using cutadapt (Martin 2011). Sequences were aligned to the reference genome release Hg38 by hla-mapper v.3.0.7, function DNA, database version 3.1, using default parameters (Castelli et al. 2018). Genotyping was performed using Toolkit’s Best Practices Workflow (GATK, v.3.7, McKenna et al. 2010). The Vcfx algorithms checkpl and evidence (www.castelli-lab.net/apps/vcfx) were used for quality control assessment so that only high-quality genotypes were maintained in subsequential analyses. The inquired genotypes were imputed and the genotypes were phased by the software phasex (available upon request), which employs Beagle v.4.1.44 (Browning and Browning 2008) to phase each VCF replicate as described in Souza et al. (2020). To create complete genomic and CDS sequences, we employed the Vcfx features “Vcfx fasta” and “Vcfx transcript” (www.castelli-lab.net/apps/vcfx), respectively. Finally, after bioinformatics quality control, 426 and 424 samples for *LILRB1* and *LILRB2* were kept for statistical analysis.

### Statistical Analysis

Expected heterozygosis, allelic and genotypic frequencies were obtained using Genepop v.4.51. Hardy-Weinberg equilibrium was tested by Guo and Thompson’s method using Arlequin v.3.5.2.

Departure from neutrality was assessed by Ewens-Watterson (Ewens 1972; Watterson 1974; Slatkin 1994), and Tajima’s *D* test (Tajima 1989) using Arlequin v.3.5.2 and Molecular Evolutionary Genetics Analysis (MEGA software version 7.0.2.1) (Kumar et al. 2016), respectively. Ewens-Watterson test compares the expected homozygosity under HWE with the expected homozygosity under the null hypothesis of neutrality-equilibrium, which enables us to test the alternative hypothesis of either directional (Observed Homozygosity > Expected Homozygosity) or balancing selection (Observed Homozygosity < Expected Homozygosity). Tajima’s *D* test explores the mean number of segregating sites (π) and the nucleotide diversity (□s), by comparing the statistics π and □s. Under neutrality, π is equal to □s, and then *D*=0. A positive *D* value (*D*>1) points out to the maintenance of different alleles in the population (heterozygous advantage), indicating balancing selection. Negative *D* values (*D*<0) point out to an excess of rare variants, with one allele more frequent than others, indicating a positive selection effect. Nonsynonymous (*d*_*N*_) and synonymous (*d*_*S*_) substitution rates were evaluated using the Nei-Gojobori method (Nei and Gojobori 1986) in MEGA v.7.0.21 (Kumar et al. 2016). Under neutrality, the *d*_*N*_/*d*_*S*_ ratio is expected to be = 1. Under purifying selection, the majority of mutations that are removed are nonsynonymous variants, resulting in *d*_*N*_/*d*_*S*_<1; while in case of positive selection, an excess of nonsynonymous fixations is expected, resulting in *d*_*N*_/*d*_*S*_>1. After coding sequences alignment by MUSCLE (Codons), Estimate Selection for each Codon (HyPhy) was used to assess the strength of selection operating upon each codon in MEGA v.7.0.21 (Kumar et al. 2016). Following sequence alignment, codon-based signatures of selection were detected using the Fast Unconstrained Bayesian Approximation (FUBAR) method (Murrell et al. 2013), implemented in the web interface Datamonkey (http://www.datamonkey.org). FUBAR approach enables the inference of nonsynonymous (dN) and synonymous (dS) substitution rates on a per-site basis for a given coding alignment and corresponding phylogeny, assuming a constant selection pressure for each site along the entire phylogeny. The approximate hierarchical Bayesian method using a Markov chain Monte Carlo (MCMC) routine employs a highly flexible and less restrictive distributional form than other random effects likelihood (REL) models, avoiding that an unrealistic constraint is applied on the distribution of selection parameters due to forced categorization of sites in a small number of classes (model misspecification). Given this distribution, FUBAR estimates the posterior probability of selection at each site in the alignment, with Bayesian posterior probabilities greater than 0.9 considered to provide enough support for positive selection (Murrell et al 2013).

Linkage disequilibrium (LD) was estimated using Haploview (Barrett et al. 2005) with haplotype blocks defined by the method of Gabriel et al. (2002), considering only variable sites with a minimum allele frequency (MAF) of 1%. Haplotype networks were inferred using the Median-Joining algorithm implemented in POPART (Population Analysis with Reticulate Trees(Leigh and Bryant 2015). Functional annotations were obtained using SnpEff (http://snpeff.sourceforge.net/) (Cingolani et al. 2012).

High coverage 1000 Genomes Project (1KGen) phase3 release data for 2,504 individuals(ftp://ftp.1000genomes.ebi.ac.uk/vol1/ftp/data_collections/1000G_2504_high_coverage/working/20190425) were downloaded considering the CDS coordinates for *LILRB1* (NM_006669.6) and *LILRB2* (NM_001080978.4), to compare the variability found in the Brazilian population with those found in the 1000 Genomes Project populations: African (AFR), European (EUR), Admixed American (AMR), East Asian (EAS), and South Asian (SAS). Thus, the genotypic data obtained from our population were compared to the 1000 Genomes data available.

### Structural analysis

Single point mutations may cause an impact on protein thermostability and dynamics. As a consequence, they may affect protein activity. To estimate the impact that these mutations may have on ILTs tertiary structures, we used an elastic network contact model to evaluate vibrational entropy changes – Elastic Network Contact Model (ENCoM; Frappier and Najmanovich 2014) and an empirical force field algorithm FoldX4 (Guerois et al. 2002; Schymkowitz et al. 2005), which calculates free energy changes between mutant LILRBs and wild type (WT).

It has been shown that the best structure-based stability predictive method corresponds to a combination of ENCoM and FoldX4 (Frappier et al. 2015). ENCoM is an entropy-based methodology to predict the effect of mutations on protein stability and dynamics (Frappier and Najmanovich 2014). As an output, ENCoM provides ΔSvib>0 as a measure of conferred rigidity and protein stability increase due to a specific mutation.

FoldX4 is an enthalpy-based approach developed for measuring the impact of mutations on the stability of proteins and nucleic acids, returning negative ΔΔG values for stabilizing mutations and positive ΔΔG values for destabilizing mutations. As previously suggested, we used 0.46 kcal/mol as a threshold for the FoldX4 accuracy (Schymkowitz et al. 2005).

LILRB1 and LILRB2 crystal structures files were downloaded from the Protein data Bank (PDB) (Berman et al. 2002): LILRB1 domains 1 and 2 (LILRB1/D1D2) (PDB 1G0X; 2.1 Å) (Chapman et al. 2000), LILRB1 domains 3 and 4 (LILRB1/D3D4) (PDB:4LL9; 2.69 Å) (Nam et al. 2013), LILRB2 domains 1 and 2 (LILRB2/D1D2) (PDB: 2DYP; 2.5 Å) (Shiroishi and Maenaka 2009) and LILRB2 domains 3 and 4 (LILRB2/D3D4) (PDB: 4LLA; 2.5 Å) (Nam et al. 2013). Single point mutations were created by Modeller 9.21 software (Sali and Blundell 1993) using PDB structures files as input. PyMol software was used for visual analysis (DeLano 2002). Predicted ΔΔG (kcal/mol) and ΔS_vib_ (kcal/mol/K) values were obtained with comparisons against the starting structure (WT).

## Results

Considering the *LILRB1* coding sequence, 58 SNVs were identified, among which one showed significant departure from Hardy-Weinberg equilibrium expectations (rs16985478, p=0.0118, Table 1). Among them, 38 had a Minor Allele Frequency (MAF) higher than 1%.

**Table 1.** Description of the 58 *LILRB1* coding sequence variants identified in an admixed Brazilian population sample.

| SNP id | GRCh38<br>Position in<br>chromosome<br>19 | *LILRB1*<br>segment | Annotation | REF | ALT | *Ho* | *He* | HWE<br>*p*-value | Alternative<br>allele<br>frequency<br>(2*n*=852) |
| --- | --- | --- | --- | --- | --- | --- | --- | --- | --- |
| rs148624633 | 54631085 | exon 3 | missense | C | G | 0.0258 | 0.0255 | 1.0000 | 0.0129 |
| rs1985501 | 54631288 | exon 4 | synonymous | C | A | 0.4953 | 0.4922 | 0.9214 | 0.5646 |
| rs61738513 | 54631627 | exon 5 | synonymous | A | T | 0.0164 | 0.0163 | 1.0000 | 0.0082 |
| rs1061679 | 54631632 | exon 5 | missense | C | T | 0.4695 | 0.4931 | 0.3312 | 0.5610 |
| rs148931844 | 54631643 | exon 5 | missense | C | T | 0.0117 | 0.0117 | 1.0000 | 0.0059 |
| rs769506618 | 54631702 | exon 5 | missense | A | T | 0.0024 | 0.0024 | 1.0000 | 0.0012 |
| rs12462774 | 54631705 | exon 5 | synonymous | C | T | 0.2676 | 0.2935 | 0.0941 | 0.8216 |
| rs12460501 | 54631706 | exon 5 | missense | A | G | 0.2676 | 0.2935 | 0.0971 | 0.8216 |
| rs147024305 | 54631712 | exon 5 | missense | C | T | 0.0024 | 0.0024 | 1.0000 | 0.0012 |
| rs61738512 | 54631736 | exon 5 | missense | G | C | 0.0258 | 0.0301 | 0.0891 | 0.0153 |
| rs138410838 | 54631748 | exon 5 | missense | C | T | 0.0352 | 0.0346 | 1.0000 | 0.0176 |
| rs10418732 | 54631750 | exon 5 | synonymous | C | G | 0.0352 | 0.0346 | 1.0000 | 0.0176 |
| rs10418552 | 54631757 | exon 5 | missense | A | C | 0.0329 | 0.0324 | 1.0000 | 0.0164 |
| rs10419085 | 54631758 | exon 5 | missense | G | C | 0.0329 | 0.0324 | 1.0000 | 0.0164 |
| rs753059498 | 54631765 | exon 5 | synonymous | C | T | 0.0024 | 0.0024 | 1.0000 | 0.0012 |
| rs778057754 | 54631770 | exon 5 | missense | T | G | 0.0024 | 0.0024 | 1.0000 | 0.0012 |
| rs1061680 | 54632001 | exon 6 | missense | C | T | 0.4531 | 0.4778 | 0.3112 | 0.6068 |
| rs1061681 | 54632040 | exon 6 | missense | T | G | 0.4648 | 0.4878 | 0.3712 | 0.5798 |
| rs2781773 | 54632064 | exon 6 | missense | A | C | 0.0024 | 0.0024 | 1.0000 | 0.0012 |
| rs182861371 | 54632235 | exon 6 | missense* | T | C | 0.0047 | 0.0047 | 1.0000 | 0.0023 |
| rs61737863 | 54632507 | exon 7 | missense | C | G | 0.0070 | 0.0070 | 1.0000 | 0.0035 |
| rs10427127 | 54632531 | exon 7 | synonymous | T | C | 0.1972 | 0.2002 | 0.8074 | 0.1127 |
| rs201906215 | 54632537 | exon 7 | synonymous | G | A | 0.0047 | 0.0047 | 1.0000 | 0.0023 |
| rs61737891 | 54632649 | exon 7 | missense | A | G | 0.0610 | 0.0592 | 1.0000 | 0.0305 |
| rs141840935 | 54632675 | exon 7 | synonymous | C | T | 0.0141 | 0.0140 | 1.0000 | 0.0070 |
| rs200461258 | 54632687 | exon 7 | stop_gained | C | G | 0.0047 | 0.0047 | 1.0000 | 0.0023 |
| rs10425827 | 54632690 | exon 7 | synonymous | A | G | 0.1455 | 0.1510 | 0.5121 | 0.0822 |
| rs74906426 | 54632696 | exon 7 | synonymous | C | T | 0.0047 | 0.0047 | 1.0000 | 0.0023 |
| rs192288587 | 54632718 | exon 7 | missense | G | A | 0.0047 | 0.0047 | 1.0000 | 0.0023 |
| rs61737895 | 54632728 | exon 7 | missense | C | T | 0.0141 | 0.0140 | 1.0000 | 0.0070 |
| rs272423 | 54632735 | exon 7 | synonymous | C | T | 0.4507 | 0.4540 | 0.9146 | 0.6526 |
| rs139074994 | 54632756 | exon 7 | missense | C | G | 0.0329 | 0.0324 | 1.0000 | 0.0164 |
| rs61739173 | 54633105 | exon 8 | missense | G | A | 0.1268 | 0.1311 | 0.4527 | 0.0704 |
| rs61739171 | 54633172 | exon 8 | missense | C | T | 0.0634 | 0.0614 | 1.0000 | 0.0317 |
| rs61739244 | 54633233 | exon 8 | synonymous | G | A | 0.0540 | 0.0526 | 1.0000 | 0.0270 |
| rs61739175 | 54633259 | exon 8 | missense | A | T | 0.0892 | 0.0853 | 1.0000 | 0.0446 |
| rs61739176 | 54633260 | exon 8 | missense | G | C | 0.0892 | 0.0853 | 1.0000 | 0.0446 |
| rs61739180 | 54633269 | exon 8 | missense | A | C | 0.0986 | 0.0941 | 0.6494 | 0.0469 |
| rs61739180 | 54633269 | exon 8 | frameshift | A | AC | 0.0986 | 0.0941 | 0.6494 | 0.0024 |
| rs200928068 | 54633296 | exon 8 | missense | C | G | 0.0070 | 0.0070 | 1.0000 | 0.0035 |
| rs2114511 | 54633642 | exon 9 | synonymous | G | C | 0.1455 | 0.1391 | 0.4960 | 0.0751 |
| rs370595875 | 54634002 | exon 10 | synonymous | C | T | 0.0094 | 0.0094 | 1.0000 | 0.0047 |
| rs1138736 | 54634003 | exon 10 | missense | G | A | 0.1338 | 0.1451 | 0.1658 | 0.0786 |
| rs1138737 | 54634655 | exon 11 | missense | C | G | 0.0751 | 0.0724 | 1.0000 | 0.0376 |
| rs1061683 | 54634688 | exon 11 | missense | A | G | 0.2183 | 0.2092 | 0.4882 | 0.1185 |
| rs10416070 | 54635114 | exon 12 | synonymous | G | A | 0.2136 | 0.2128 | 1.0000 | 0.1209 |
| rs61737692 | 54635585 | exon 14 | synonymous | G | A | 0.0211 | 0.0209 | 1.0000 | 0.0106 |
| rs113759647 | 54635591 | exon 14 | synonymous | G | T | 0.0117 | 0.0117 | 1.0000 | 0.0059 |
| rs200167361 | 54636520 | exon 15 | missense | G | C | 0.0047 | 0.0047 | 1.0000 | 0.0023 |
| rs41308744 | 54636536 | exon 15 | missense | G | C | 0.0657 | 0.0636 | 1.0000 | 0.0329 |
| rs202204734 | 54636537 | exon 15 | missense | A | C | 0.0657 | 0.0636 | 1.0000 | 0.0329 |
| rs200490901 | 54636553 | exon 15 | missense | A | T | 0.0164 | 0.0163 | 1.0000 | 0.0082 |
| rs41308746 | 54636580 | exon 15 | synonymous | T | C | 0.2324 | 0.2304 | 1.0000 | 0.1326 |
| rs28414191 | 54636592 | exon 15 | synonymous | T | C | 0.1620 | 0.1723 | 0.2513 | 0.0951 |
| rs28409473 | 54636594 | exon 15 | missense | G | A | 0.1596 | 0.1665 | 0.3789 | 0.0916 |
| rs62133433 | 54636625 | exon 15 | synonymous | G | A | 0.1620 | 0.1723 | 0.2515 | 0.0951 |
| rs1061684 | 54636791 | exon 16 | synonymous | C | T | 0.2676 | 0.2457 | 0.0743 | 0.1432 |
| rs16985478 | 54636798 | exon 16 | missense | G | A | 0.2418 | 0.2164 | **0.0118** | 0.1232 |
| rs41548213 | 54636848 | exon 16 | synonymous | C | A | 0.0634 | 0.0614 | 1.0000 | 0.0317 |

\* Variant may also be within a splice site
\*\* HWE *p*-value below 0.05 was considered significant and highlighted in bold.

These *LILRB1* 58 SNVs configure 84 haplotypes, 13 of which with frequency higher than 1% in these Brazilian population sample (Table S1). The haplotypes were named sequentially according to their frequencies within each of the three haplotype groups defined in the network in Figure 1. Variation sites shared by a given haplotype group were identified and highlighted in Table S1. Three major haplotypes accounted for almost two thirds (64.8%) of the haplotype variation: a1 (37.6%), b1 (16.7%), and b2 (10.6%).

**Figure 1.**
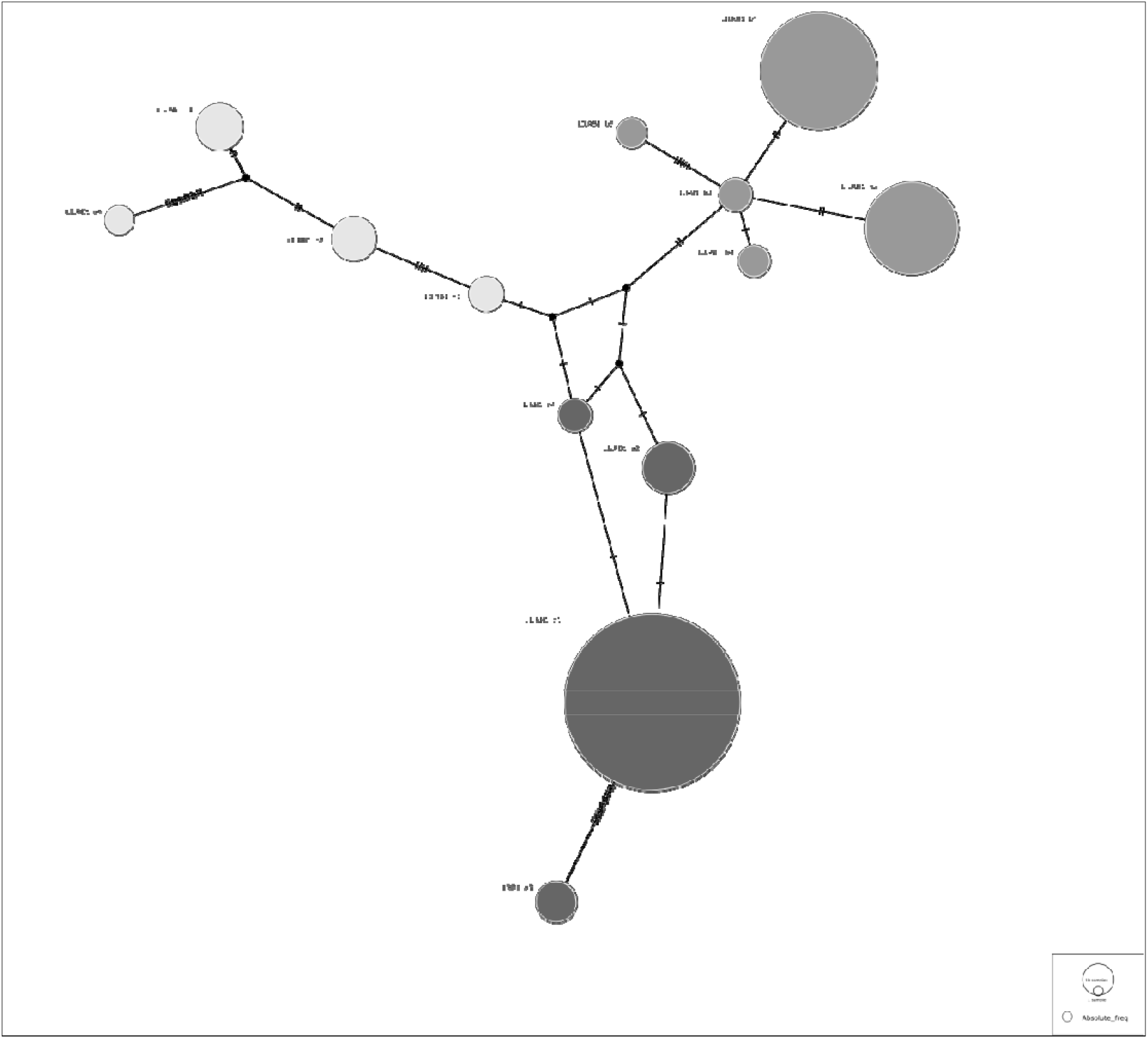
Haplotype network illustrating the similarity among the *LILRB1* haplotypes having a frequency > 1% in the Brazilian population. Haplotype groups and members inside each haplotype group were named considering their frequencies in descending order.

Linkage disequilibrium was also assessed along the *LILRB1* segment, and the LD pattern was observed with four segregation blocks spread across the entire gene region (Figure 2).

**Figure 2.**
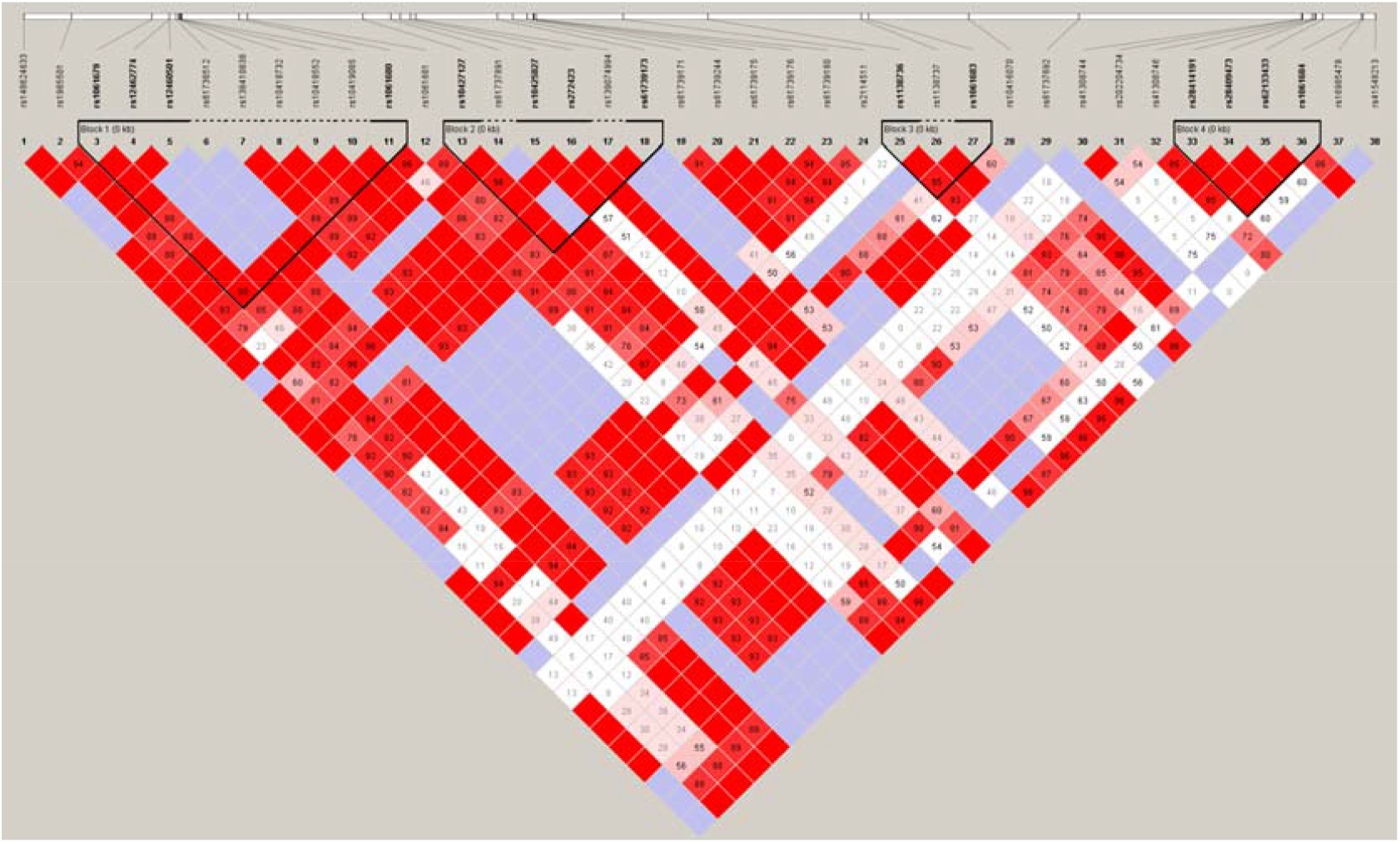
Linkage disequilibrium among the SNPs from coding region of LILRB1. The first LD block comprised variants from exons 5 and 6, the second block included variants from exons 7 and 8, the third block, variants from exons 10 and 11, and the fourth block, variants from exons 15 and 16. LD is evaluated by D’ values and LOD scores. Tagger SNPs are highlighted in bold. Areas in red indicate strong LD (LOD ≥ 2, D’ = 1); areas in light red indicate moderate LD (LOD ≥ 2, D’ < 1); areas in blue indicate LD with lack of statistical evidence (LOD ≤ 2, D’ = 1); while areas in white indicate no LD (LOD ≤ 2, D’ < 1). D’ values different from 1.00 are represented inside the squares as percentages. LOD, log of the odds; D’, pairwise correlation between single-nucleotide polymorphisms.

Regarding *LILRB2* coding regions, 41 SNVs were identified in the Brazilian sample evaluated in this study, three of them deviating from the Hardy-Weinberg equilibrium expectations. Almost half of them (20 out of 41) had a MAF higher than 1% (Table 2).

**Table 2.** Description of the 41 variation sites identified in *LILRB2* coding sequence in an admixed Brazilian population sample.

| SNP id | GRCh38<br>position in<br>chromosome<br>19 | *LILRB2*<br>segment | Annotation | REF | ALT | *Ho* | *He* | HWE<br>*p*-<br>value | Alternative<br>allele<br>frequency<br>(2*n*=848) |
| --- | --- | --- | --- | --- | --- | --- | --- | --- | --- |
| rs144888744 | 54274698 | exon 14 | synonymous | G | A | 0.0024 | 0.0024 | 1.0000 | 0.0012 |
| rs143169595 | 54274730 | exon 14 | missense | C | T | 0.0047 | 0.0047 | 1.0000 | 0.0024 |
| rs146683137 | 54274733 | exon 14 | missense | T | C | 0.0330 | 0.0325 | 1.0000 | 0.0165 |
| rs143326661 | 54274763 | exon 14 | missense | T | C | 0.0024 | 0.0024 | 1.0000 | 0.0012 |
| rs151307136 | 54274770 | exon 14 | synonymous | G | A | 0.0047 | 0.0047 | 1.0000 | 0.0024 |
| rs144872171 | 54274812 | exon 14 | synonymous | G | C | 0.0071 | 0.0071 | 1.0000 | 0.0035 |
| rs35440540 | 54275952 | exon 13 | missense* | C | T | 0.0094 | 0.0094 | 1.0000 | 0.0047 |
| rs201537103 | 54275975 | exon 13 | synonymous | T | C | 0.0283 | 0.0279 | 1.0000 | 0.0142 |
| rs149508154 | 54276808 | exon 10 | synonymous* | C | T | 0.0283 | 0.0279 | 1.0000 | 0.0142 |
| rs61063260 | 54277554 | exon 09 | synonymous | T | C | 0.0071 | 0.0071 | 1.0000 | 0.0035 |
| rs368934809 | 54277563 | exon 09 | synonymous | C | T | 0.0047 | 0.0047 | 1.0000 | 0.0024 |
| rs143311942 | 54277908 | exon 08 | synonymous | G | A | 0.0024 | 0.0024 | 1.0000 | 0.0012 |
| rs4993133 | 54278311 | exon 07 | missense | C | T | 0.0024 | 0.0024 | 1.0000 | 0.0012 |
| rs142656214 | 54278395 | exon 07 | missense | G | C | 0.0024 | 0.0024 | 1.0000 | 0.0012 |
| rs759172779 | 54278402 | exon 07 | synonymous | G | A | 0.0024 | 0.0024 | 1.0000 | 0.0012 |
| rs138762759 | 54278436 | exon 07 | missense | G | A | 0.0047 | 0.0047 | 1.0000 | 0.0024 |
| rs149499521 | 54278469 | exon 07 | missense | T | G | 0.0071 | 0.0071 | 1.0000 | 0.0035 |
| rs7247025 | 54278473 | exon 07 | missense | G | A | 0.1722 | 0.1922 | **0.0247** | 0.0118 |
| rs7247025 | 54278473 | exon 07 | missense | G | C | 0.1722 | 0.1922 | **0.0247** | 0.0943 |
| rs114686155 | 54278525 | exon 07 | synonymous | T | C | 0.0189 | 0.0187 | 1.0000 | 0.0094 |
| rs28405793 | 54278534 | exon 07 | synonymous | T | C | 0.1156 | 0.1256 | 0.1098 | 0.0672 |
| rs116027944 | 54278539 | exon 07 | missense | T | G | 0.1179 | 0.1276 | 0.1199 | 0.0684 |
| rs7246737 | 54278541 | exon 07 | missense | A | G | 0.1179 | 0.1276 | 0.1212 | 0.0684 |
| rs7247055 | 54278547 | exon 07 | missense | G | C | 0.1156 | 0.1296 | **0.0429** | 0.0696 |
| rs1128646 | 54278553 | exon 07 | missense | C | T | 0.4717 | 0.4952 | 0.3756 | 0.4481 |
| rs10405713 | 54278815 | exon 06 | missense | T | C | 0.0684 | 0.0749 | 0.1252 | 0.0389 |
| rs41308754 | 54278816 | exon 06 | missense | G | C | 0.0330 | 0.0325 | 1.0000 | 0.0165 |
| rs150523334 | 54278817 | exon 06 | missense | A | G | 0.0024 | 0.0024 | 1.0000 | 0.0012 |
| rs7247451 | 54278849 | exon 06 | missense | G | C | 0.4623 | 0.4981 | 0.1398 | 0.4646 |
| rs7247208 | 54278855 | exon 06 | synonymous | A | G | 0.4788 | 0.4991 | 0.4360 | 0.4729 |
| rs7247538 | 54278869 | exon 06 | missense | G | A | 0.4670 | 0.4961 | 0.2403 | 0.4528 |
| rs10404685 | 54278876 | exon 06 | synonymous | G | A | 0.1085 | 0.1235 | **0.0267** | 0.0660 |
| rs41308750 | 54278882 | exon 06 | synonymous | T | C | 0.0354 | 0.0348 | 1.0000 | 0.0177 |
| rs143661416 | 54278899 | exon 06 | missense | A | G | 0.0047 | 0.0047 | 1.0000 | 0.0024 |
| rs386056 | 54279064 | exon 06 | missense | T | C | 0.2429 | 0.2680 | 0.0687 | 0.8408 |
| rs143879306 | 54279460 | exon 05 | synonymous | G | A | 0.0094 | 0.0094 | 1.0000 | 0.0047 |
| rs373032 | 54279520 | exon 05 | missense | A | T | 0.3467 | 0.3442 | 1.0000 | 0.2205 |
| rs73055442 | 54279838 | exon 04 | missense | C | T | 0.0519 | 0.0506 | 1.0000 | 0.0259 |
| rs111753747 | 54279942 | exon 04 | synonymous | A | G | 0.0024 | 0.0024 | 1.0000 | 0.0012 |
| rs366337 | 54280068 | exon 04 | synonymous | G | A | 0.1274 | 0.1235 | 1.0000 | 0.0660 |
| rs45557835 | 54280267 | exon 03 | missense | T | C | 0.0024 | 0.0024 | 1.0000 | 0.0012 |
| rs383369 | 54280275 | exon 03 | missense | G | A | 0.2288 | 0.2517 | 0.0780 | 0.8526 |

\* Variant may also be within a splice site
\*\* HWE *p*-values below 0.05 were considered significant and highlighted in bold.

A total of 55 *LILRB2* haplotypes were inferred from these 41 SNVs, 11 of which occurred at a frequency higher than 1% in this admixed Brazilian population sample (Table S2). These 11 haplotypes were named sequentially according to their frequencies within each of the four haplotype groups (Figure 3). Variation sites shared by a given haplotype group were identified and highlighted in Table S2. As observed for *LILRB2*, three haplotypes were predominant, accounting for almost two thirds (63.3%): a1 (36.7%), b1 (13.2%), and c1 (13.4%).

**Figure 3.**
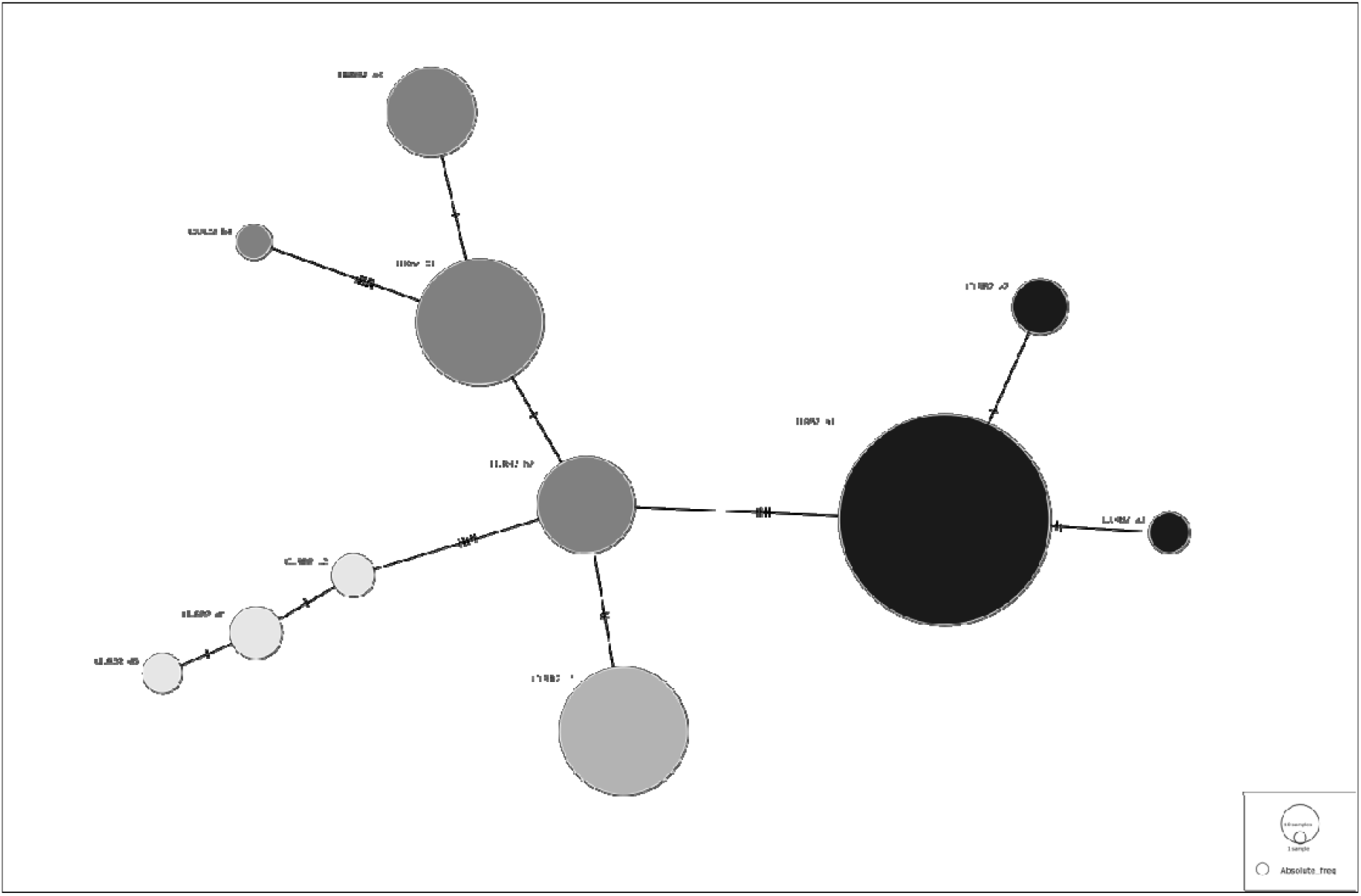
Haplotype network illustrating the sequence similarity among the 11 *LILRB2* haplotypes occurring at a frequency of at least 1% in the Brazilian population. Haplotype groups and members inside each haplotype group were named considering their frequencies in descending order.

Linkage disequilibrium patterns were also assessed along the *LILRB2* segment (Figure 4), and the LD pattern demonstrated the presence of four small segregation blocks concentrated in the initial gene segment (from exon 3 to exon 7).

**Figure. 4.**
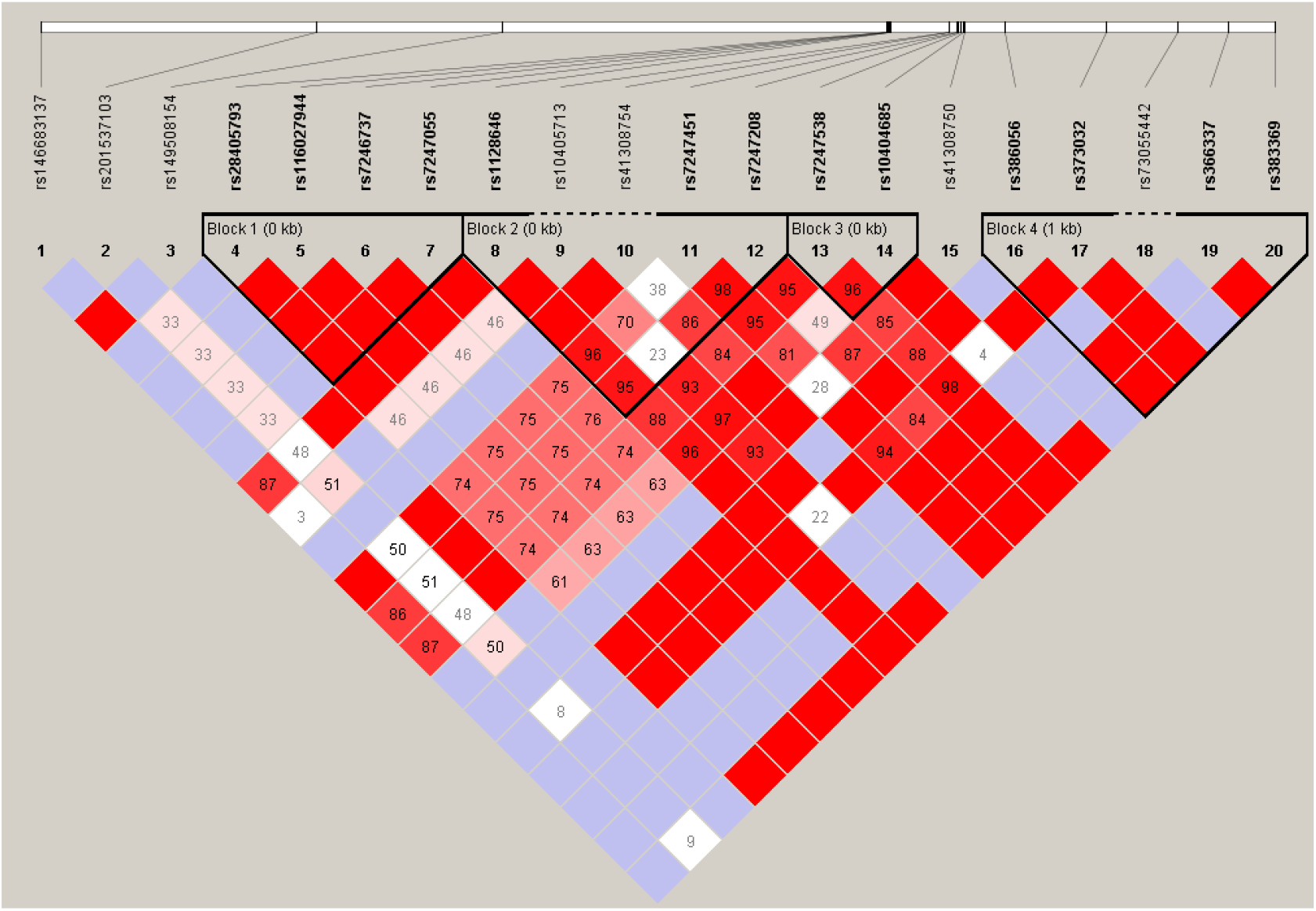
Linkage disequilibrium among the SNPs from coding region of *LILRB2*. The first LD block comprises only variants from exon 7, the second block has variants from exons 6 and 7, the third block contains only variants from exon 6, and meanwhile the fourth block 4 comprises variants from exons 3-6. LD is evaluated by *D’* values and *LOD* scores. Tagger SNPs are highlighted in bold. Areas in red indicate strong LD (*LOD* ≥ 2, *D’* = 1); areas in light red indicate moderate LD (*LOD* ≥ 2, *D’* < 1); areas in blue indicate LD with lack of statistical evidence (*LOD* ≤ 2, *D’* = 1); while areas in white indicate no LD (*LOD* ≤ 2, *D’* < 1). *D’* values different from 1.00 are represented inside the squares as percentages. *LOD*, log of the odds; *D’*, pairwise correlation between single-nucleotide polymorphisms.

To evaluate whether natural selection may have influenced allele and genotype frequencies, we applied four different neutrality tests on the genotype data of the population sample under study. For the majority of tests, the coding regions of each gene were subdivided into groups of exons according to the minimum number of 10 variants required by the software MEGAv.7.0.2.1 (Table 3), while the analysis using FUBAR considered the whole coding region under investigation.

**Table 3.** Results of Tajima’s *D* and Ewens-Watterson neutrality test. *P*-values considered significant in two-tailed Tajima’s *D* (*p*< 0.025) and Ewens-Watterson (*p*> 0.975) neutrality tests are highlighted in bold.

| Neutrality tests | Tajima's *D* | Ewens–Watterson's neutrality test |  |  |
| --- | --- | --- | --- | --- |
|  |  | *F* observed | *F* expected | *p*-value |
| *LILRB1* |  |  |  |  |
| entire gene | *D* = - **0.2310**; *P* = **0.0018** | 0.1845 | 0.0431 | *P* = **1.0000** |
| exons 3, 4, 5 | *D* = - **1.0470**; *P* = **0.0055** | 0.3641 | 0.2753 | *P* = 0.9370 |
| exons 6, 7 | *D* = - **0.7629**; *P* = **0.0047** | 0.3730 | 0.2319 | *P* = **0.9860** |
| exons 8, 9, 10 | *D* = **1.0735**; *P* = **0.0044** | 0.7014 | 0.2814 | *P* = 0.9640 |
| exons 11, 12, 14, 15, 16 | *D* = **0.9298**; *P* = **0.0023** | 0.4684 | 0.1302 | *P* = **0.9970** |
| *LILRB2* |  |  |  |  |
| entire gene | *D* = - **1.0723**; *P* = **0.0011** | 0.1828 | 0.0717 | *P* = **1.0000** |
| exons 3, 4, 5, 6 | *D* = - **0.1532**; *P* = **0.0030** | 0.2239 | 0.1903 | *P* = 0.9020 |
| exon 7 | *D* = - **0.4374**; *P* = **0.0135** | 0.3909 | 0.2792 | *P* = **0.9910** |
| exons 8, 9, 10, 13, 14 | *D* = - **1.8303**; *P* = **0.0007** | 0.9010 | 0.3319 | *P* = **1.0000** |

Results of the Ewens-Watterson test for neutrality for *LILRB1* showed statistically significant positive normalized *F* values (observed homozygosity significantly higher than expected homozygosity, with *p*-values above 0.975) for the whole gene as well as for all exons except exons 3, 4 and 5, and 8, 9 and 10. Similar results were observed for *LILRB2*, with significant positive normalized *F* values for the whole gene and all exons except exons 3, 4, 5, 6 (Table 3). Tajima’s *D* test showed statistically significant negative *D* values for three segments of *LILRB1* (whole gene, exons 3, 4, 5 and exons 6, 7) and for the four *LILRB2* regions evaluated (Table 3). Nonetheless, significantly positive *D* values were observed for the segments encompassing exons 8, 9, 10 and 11, 12, 14, 15, 16 of *LILRB1*.

The synonymous and nonsynonymous nucleotide substitution test enables to evaluate hypotheses of either purifying or positive selection, with *d*_*N*_/*d*_*S*_ values higher than 1, lower than 1, and equal to 1 indicating positive selection, purifying selection, and neutral evolution, respectively (Table 4). Considering *LILRB1*, we observed no significant *d*_*N*_/*d*_*S*_ results, whereas a single signal of positive selection was observed for *LILRB2* (segment involving exons 3, 4, 5, 6).

**Table 4.** Results of *d*_*N*_/*d*_*S*_ test for *LILRB1* and *LILRB2* coding regions. Significant *p*-values are highlighted in bold.

| Neutrality tests | Number of sequences | $H_A$=Neutrality ($d_N \neq d_S$) | $H_A$=Positive Selection ($d_N > d_S$) | $H_A$=Purifying Selection ($d_N < d_S$) |
| --- | --- | --- | --- | --- |
| **LILRB1** |  |  |  |  |
| entire gene | 84 | $dN - dS = 0.3070$; $P = 0.7590$ | $dN - dS = 0.3030$; $P = 0.3810$ | $dN - dS = 0.3070$; $P = 1.0000$ |
| exons 3, 4, 5 | 14 | $dN - dS = 0.4790$; $P = 0.6330$ | $dN - dS = 0.4910$; $P = 1.0000$ | $dN - dS = 0.5240$; $P = 0.3010$ |
| exons 6, 7 | 20 | $dN - dS = 1.2840$; $P = 0.2020$ | $dN - dS = 1.3280$; $P = 1.0000$ | $dN - dS = 1.2910$; $P = 0.1000$ |
| exons 8, 9, 10 | 15 | $dN - dS = 0.2090$; $P = 0.8350$ | $dN - dS = 0.2060$; $P = 1.0000$ | $dN - dS = 0.2090$; $P = 0.4170$ |
| exons 11, 12, 14, 15, 16 | 32 | $dN - dS = 0.3740$; $P = 0.7090$ | $dN - dS = 0.3610$; $P = 1.0000$ | $dN - dS = 0.3610$; $P = 0.3590$ |
| **LILRB2** |  |  |  |  |
| entire gene | 55 | $dN - dS = 0.5070$; $P = 0.6130$ | $dN - dS = 0.5380$; $P = 1.0000$ | $dN - dS = 0.5140$; $P = 0.3040$ |
| exons 3, 4, 5, 6 | 23 | $dN - dS = 1.9415$; $P = 0.0545$ | $dN - dS = 1.9165$; $P = 0.0288$ | $dN - dS = -1.9380$; $P = 1.0000$ |
| exon 7 | 15 | $dN - dS = -0.9110$; $P = 0.3640$ | $dN - dS = -9.1000$; $P = 1.0000$ | $dN - dS = 0.9020$; $P = 0.1850$ |
| exons 8, 9, 10, 13, 14 | 12 | $dN - dS = -1.5470$; $P = 0.1250$ | $dN - dS = -1.5730$; $P = 1.0000$ | $dN - dS = 1.5899$; $P = 0.0572$ |

Lastly, we used the Bayesian approach implemented in FUBAR to evaluate nonsynonymous and synonymous substitution rates on a per-site basis. FUBAR identified nine codons under positive selection and seven codons under negative selection for *LILRB1* (Table 5), as well as five codons under positive selection and four codons under negative selection for *LILRB2* (Table 6).

**Table 5.** Information retrieved from SnpEff to *LILRB1* sites identified as a target by positive or negative selection using FUBAR.

| residue | DNA position | ID number | exon | variant notation at DNA level | natural selection effect | variant notation at protein level | Protein region |
| --- | --- | --- | --- | --- | --- | --- | --- |
| 68 | 19: 54631632 | rs1061679 | 5 | missense | positive selection | p.Pro68Leu | D1 |
| 92 | 19: 54631705 | rs12462774 | 5 | synonymous | negative selection | p.His92His | D1 |
| 142 | 19: 54632001 | rs1061680 | 6 | missense | positive selection | p.Thr142Ile | D2 |
| 155 | 19: 54632040 | rs1061681 | 6 | missense | positive selection | p.Ile155Ser | D2 |
| 298 | 19: 54632696 | rs74906426 | 7 | synonymous | negative selection | p.Tyr298Tyr | D3 |
| 392 | 19: 54633233 | rs61739244 | 8 | synonymous | negative selection | p.Ala392Ala | D4 |
| 422 | 19: 54633642 | rs2114511 | 9 | synonymous | negative selection | p.Pro422Pro | stem |
| 448 | 19: 54634003 | rs1138736 | 10 | missense | positive selection | p.Gly448Arg | stem |
| 470 | 19: 54634688 | rs1061683 | 11 | missense | positive selection | p.Ile470Val | transmembrane |
| 564 | 19: 54636536 | rs41308744 | 15 | missense | positive selection | p.Glu564Lys/Gln | ITIM motif 2 |
| 564 | 19: 54636537 | rs202204734 | 15 | missense | positive selection | p.Glu564Ala | ITIM motif 2 |
| 569 | 19: 54636553 | rs200490901 | 15 | missense | negative selection | p.Arg569Ser | cytoplasmatic tail |
| 582 | 19: 54636592 | rs28414191 | 15 | synonymous | negative selection | p.Ser582Ser | cytoplasmatic tail |
| 583 | 19: 54636594 | rs28409473 | 15 | missense | positive selection | p.Gly583Glu | cytoplasmatic tail |
| 593 | 19: 54636625 | rs62133433 | 15 | synonymous | negative selection | p.Ala593Ala | cytoplasmatic tail |
| 625 | 19: 54636798 | rs16985478 | 16 | missense | positive selection | p.Glu625Lys | cytoplasmatic tail |
| 641 | 19: 54636848 | rs41548213 | 16 | synonymous | negative selection | p.Pro641Pro | cytoplasmatic tail |

**Table 6.** Information retrieved from SnpEff to *LILRB2* sites identified as a target by positive or negative selection using FUBAR.

| residue | DNA position | ID number | exon | variant notation at DNA level | natural selection effect | variant notation at protein level | protein region |
| --- | --- | --- | --- | --- | --- | --- | --- |
| 297 | 19: 54278876 | rs10404685 | 6 | synonymous | negative selection | p.Tyr297Tyr | D3 |
| 300 | 19: 54278869 | rs7247538 | 6 | missense | positive selection | p.His300Tyr | D3 |
| 304 | 19: 54278855 | rs7247208 | 6 | synonymous | negative selection | p.Ser304Ser | D3 |
| 306 | 19: 54278849 | rs7247451 | 6 | missense | positive selection | p.Cys306Trp | D3 |
| 318 | 19: 54278815 | rs10405713 | 6 | missense | positive selection | p.Thr318Ala | D3 |
| 322 | 19: 54278553 | rs1128646 | 7 | missense | positive selection | p.Arg322His | D4 |
| 328 | 19: 54278534 | rs28405793 | 7 | synonymous | negative selection | p.Ser328Ser | D4 |
| 349 | 19: 54278473 | rs7247025 | 7 | missense | positive selection | p.Arg349Trp | D4 |
| 541 | 19: 54275975 | rs201537103 | 13 | synonymous | negative selection | p.Glu541Glu | cytoplasmatic tail |

We used a bioinformatics approach to evaluate the functional impact of missense substitutions. A total of 23 *LILRB1* SNVs (12 of them located in D1D2 domains and 11 in D3D4 domains) and 21 *LILRB2* SNVs (1 located in D1D2 domains and 20 in D3D4 domains) were evaluated. Concerning structural analysis, free energy changes were analyzed by FoldX4, which predicts protein stability changes (ΔΔG values in kcal/mol) due to mutations. As shown in Figure 5, FoldX4 predicted highly destabilizing changes for five missense substitutions in LILRB1 (A93T, L114R, I155S, L197P, and G350R) and four such substitutions (G317T, F326S, P375A, and P375T) in LILRB2 compared to WT (ΔΔG > 1.84 kcal/mol).

**Figure. 5.**
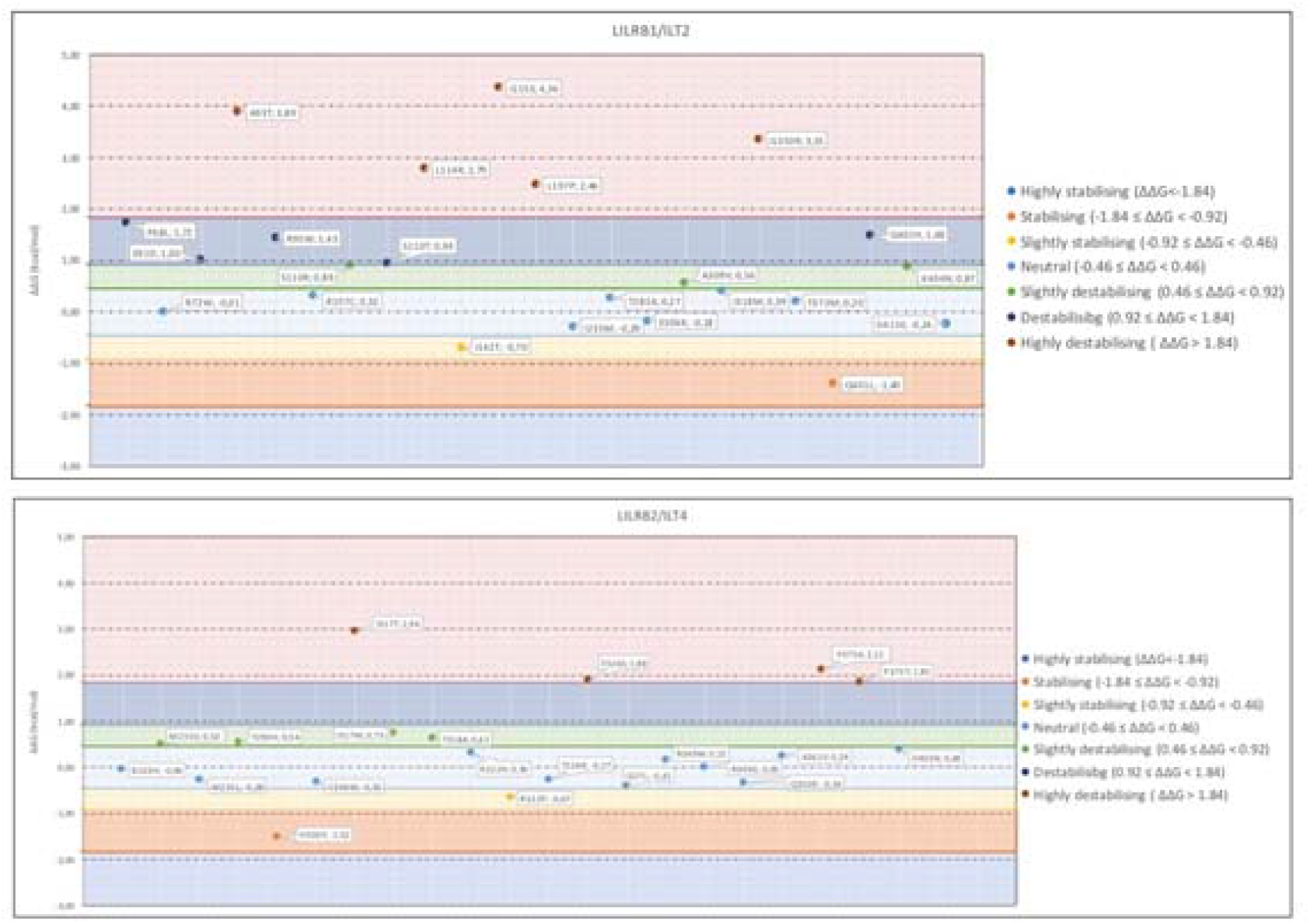
ΔΔG values (kcal/mol) for *LILRB1* and *LILRB2* variations. Altogether, five and four mutations were identified as highly destabilizing in LILRB1 (top) and LILRB2 (bottom) structures, respectively.

FoldX does not explore protein backbone conformational changes. We were also interested in evaluating if the single mutations could affect the flexibility of the entire protein when compared to the wild structure. Therefore, vibrational entropy differences (ΔSVib) (kcal/mol/Kelvin) between mutations and WT were analyzed by ENCoM. This software predicts the conformational changes caused by the exchange of protein residues comparing the new and the wild-type structures. The smaller ΔS_Vib_ the greater the flexibility and the smaller the contribution to protein stability. As a result, we observed that vibrational entropy is lower on average for LILRB1 than LILRB2 (ΔS_Vib_< 0), suggesting that *LILRB1* mutations lead to a more flexible protein (Figure 6).

**Figure 6.**
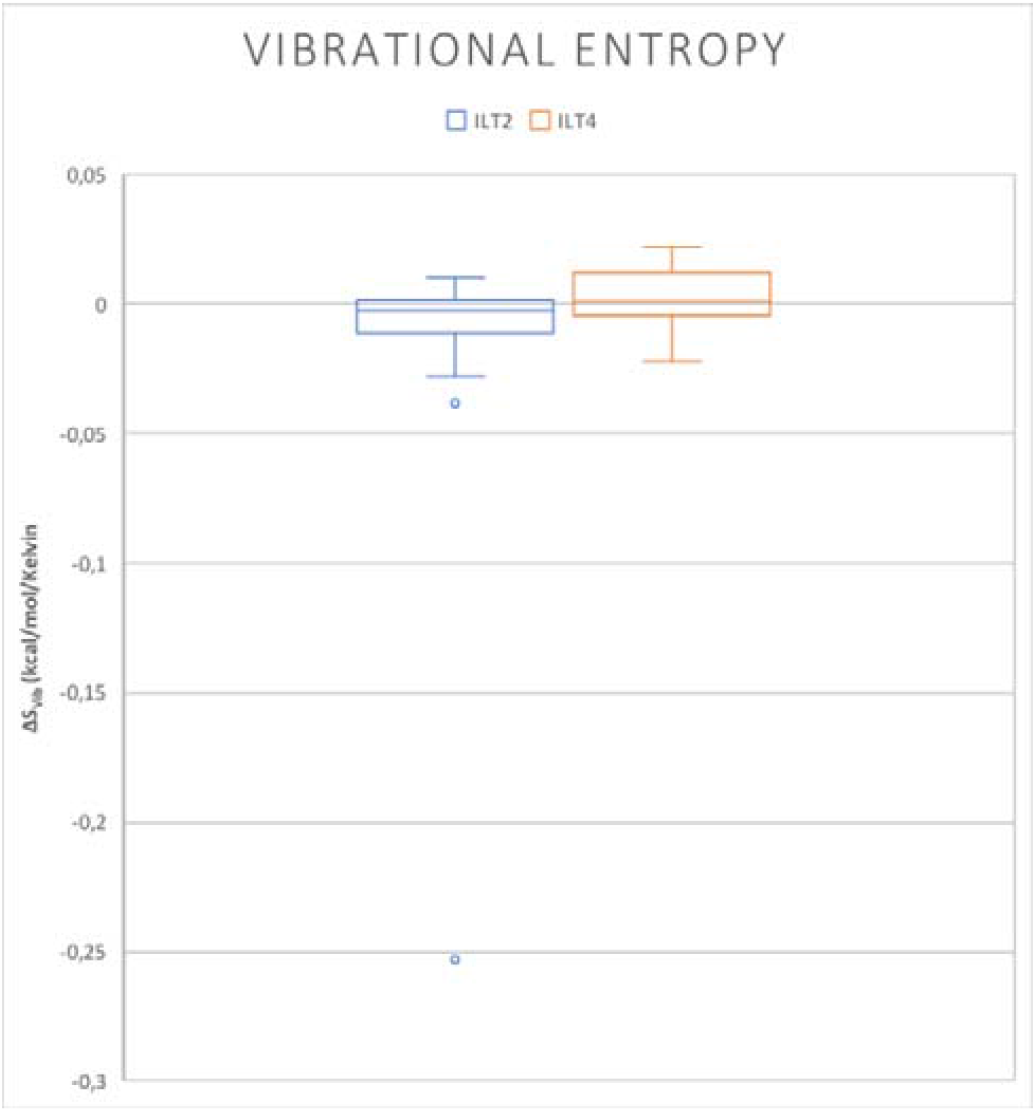
Average ΔS_Vib_ (kcal/mol). Mean difference of vibrational entropy between mutant and wild-type structures in LILRB1 (ILT2) and LILRB2 (ILT4). ΔS_Vib_> 0, less flexible and more stable; ΔS_Vib_< 0, gain of flexibility.

We also analyzed ΔS_Vib_ values of the five most affected residues for each mutation in LILRB1 (Figures 7 and 8) and LILRB2 (Figures 9 and 10). 3D structure images were classified under a color scale from blue (less flexible) to red (gain of flexibility). For each set of mutations, color values are set in terms of 3 times the standard deviation or according to the maximum absolute difference, whichever is smaller, and heatmap color scales were determined considering the maximum and minimum values of each entire set of mutations.

**Figure. 7.**
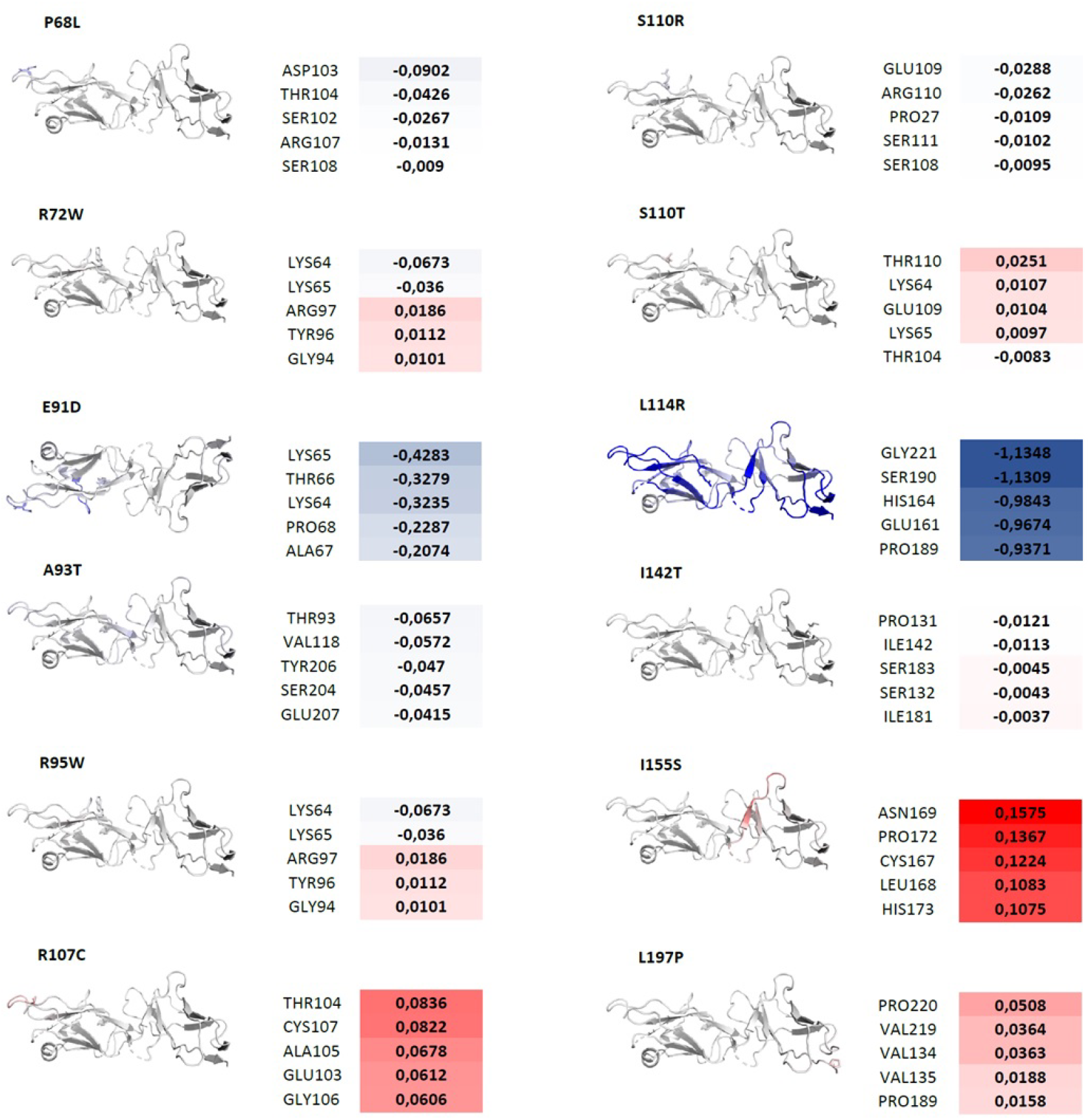
Vibrational entropy differences (ΔS_Vib_) (kcal/mol/Kelvin). ΔS_Vib_ values for the five most affected residues for the set of mutations in LILRB1/D1D2 domains. Values highlighted in blue indicate less of flexibility, while those highlighted in red indicate gain of flexibility (less stable). Maximum absolute difference = 1.1348; SD = 0.3048.

**Figure. 8.**
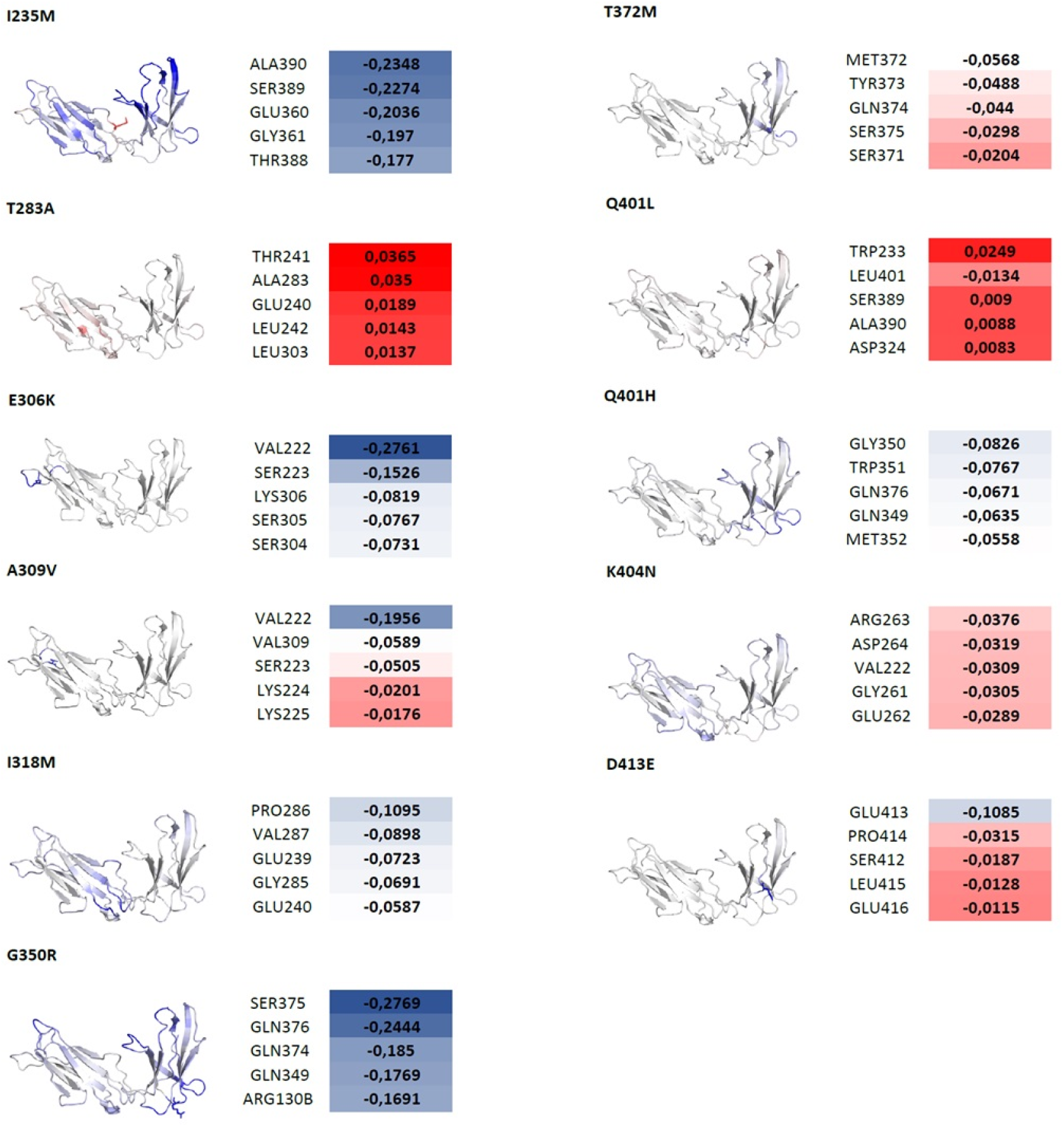
Vibrational entropy differences (ΔS_Vib_) (kcal/mol/Kelvin). ΔS_Vib_ values for the five most affected residues for the set of mutations in LILRB1/D3D4 domains. Values highlighted in blue indicate less of flexibility, while those highlighted in red indicate gain of flexibility (less stable). Maximum absolute difference = 0.2769 SD = 0.0737.

**Figure. 9.**
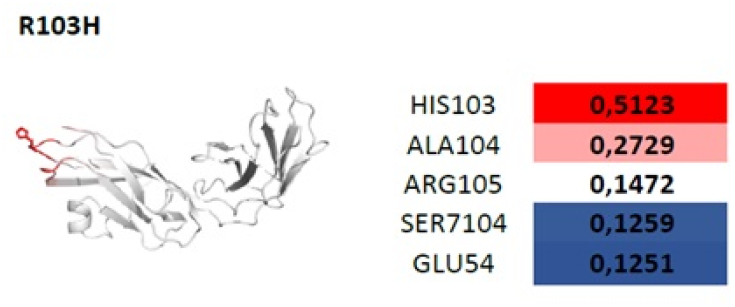
Vibrational entropy differences (ΔS_Vib_) (kcal/mol/Kelvin). ΔS_Vib_ values for the five most affected residues for the set of mutations in LILRB2/D1D2 domains. Values highlighted in blue indicate less of flexibility, while those highlighted in red indicate gain of flexibility (less stable). Maximum absolute difference = 0.5123 SD = 0.1393.

**Figure. 10.**
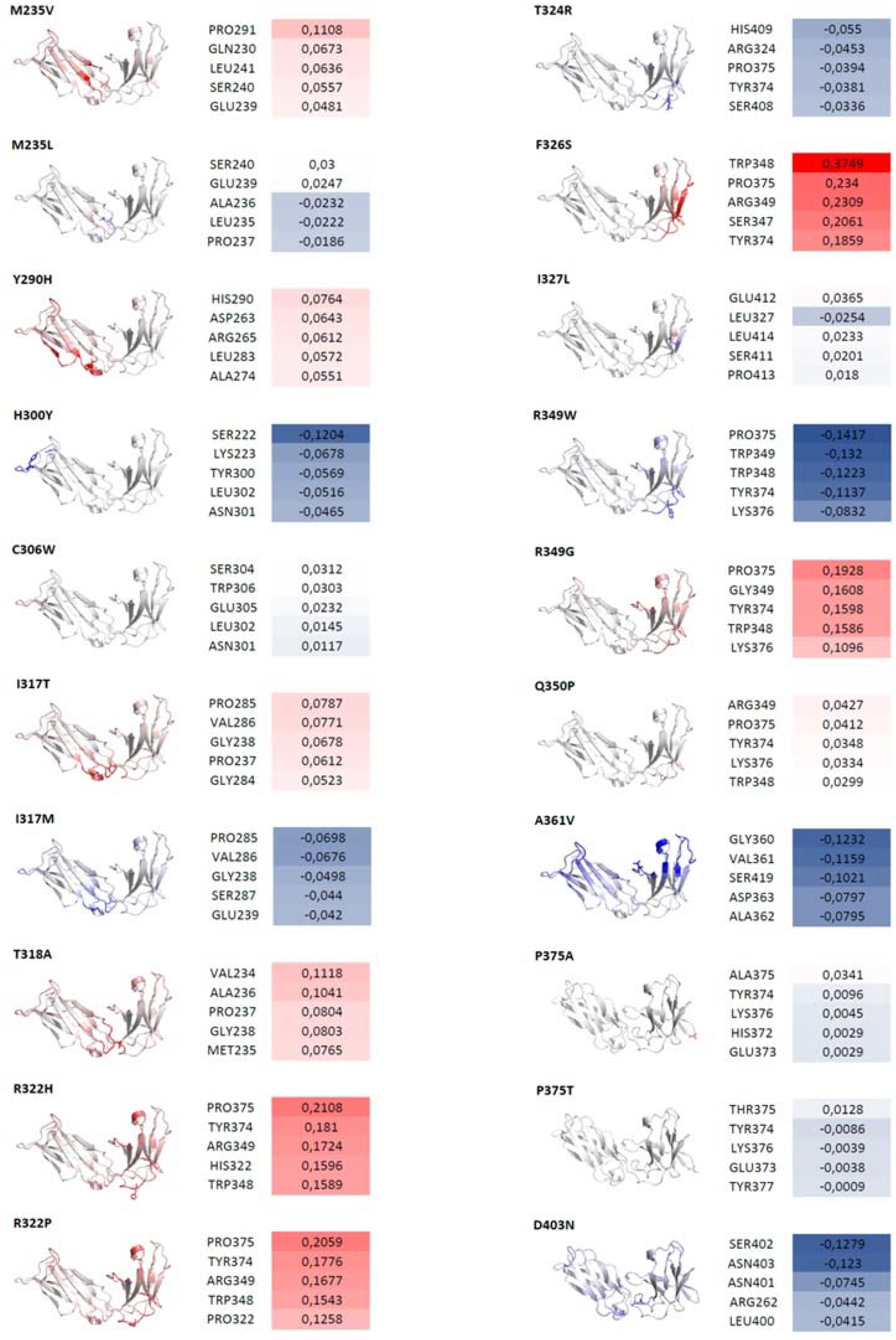
Vibrational entropy differences (ΔS_Vib_) (kcal/mol/Kelvin). ΔS_Vib_ values for the five most affected residues for the set of mutations in LILRB2/D3D4 domains. Values highlighted in blue indicate less of flexibility, while those highlighted in red indicate gain of flexibility (less stable). Maximum absolute difference = 0.3749; SD = 0.0598.

## Discussion

To the best of our knowledge, this is the first study to fully characterize the variability in all *LILRB1* and *LILRB2* exons using massively parallel sequencing. The bioinformatics strategy used here, based on a previously described methodology (Castelli et al. 2008; Souza et al. 2020), recently implemented in the hla-mapper software, allowed achieving a reliable read mapping of *LILRBs* and *KIRs* genes. The mapping step is an important issue especially for genes as *LILRs* that arise from intergenic recombination and gene duplication (Canavez et al. 2001). These genes present a high sequence similarity level between each other, with identity levels among the *LILRs* D1-D4 ranging from 63% to 84% when aligned to *LILRB1* (Borges and Cosman 2000), which may seriously bias the mapping.

Admixed populations usually harbor genomes that originated from different continental ones that have diverged because of genetic drift and diverse selective pressures (Seldin et al. 2011). This increased population diversity, with unique patterns of allelic and genotypic frequencies, linkage disequilibrium and epistatic interactions, provides a powerful approach for identifying new genetic variants, which would have been missed by exclusively studying ancestrally homogeneous populations (Ardlie et al. 2002; Rosenberg et al. 2010; Seldin et al. 2011). As a highly admixed population, the Brazilian genetic architecture makes it an ideal material for exploratory studies leading to the identification of undescribed variants (Salzano 2004). Thus, combining a NGS method, which increases the sensitivity for detection of new variants, and a bioinformatics approach to minimize mapping errors to a highly heterogeneous population, we surveyed the *LILRB1* and *LILRB2* coding regions provide further insights into the genetic diversity pattern of these essential immune regulators.

### *LILRB1* and *LILRB2* genetic diversity and LD patterns in Brazil

We have identified 58 SNVs encompassing all *LILRB1* exons (Table 1). Many of these variable sites (65.5%) show MAF higher than 1% in this Brazilian sample, and 58.6% of them correspond to missense variants, with one variant not attending HWE (rs16985478; *p*-value= 0.0118). The combination of these 58 variable sites gives rise to 84 haplotypes, 13 of them occurring with a frequency above 1% in this Brazilian population sample (Table S1). The *LILRB1* haplotype network (Figure 1) reveals the presence of 3 haplotype groups, which are defined by the missense variant rs1061679 (haplogroup a), the missense variant rs1061680 (haplogroup b) and the synonymous variant rs10416070 (haplogroup c). It is noteworthy that haplogroups a1 and b1 correspond to more than 50% of the identified haplotypes in our population (Table S1), with the SNPs that differentiate these haplogroups (rs1061679 and rs1061680) being part of the same LD block (Figure 2). Additionally, they correspond to sites located in D1/D2 identified as being under positive selection according to FUBAR and classified by FoldX4 as destabilizing mutations. In comparison, haplogroup b1 corresponds to the single haplotype harboring the reference allele in the site corresponding to rs12460501, which is interesting given that A93T (located in the D1 domain) was identified by FoldX4 analysis as being a highly destabilizing mutation.

Among the 58 SNVs identified in *LILRB1*, 25 SNPs have already been described in a previous study (Davidson et al. 2010). Others studies have also identified and studied some of these variants: rs1061680 (Kuroki et al. 2005; Bylińska et al. 2018; Yu et al. 2018), rs1061681 (Kuroki et al. 2005; Yu et al. 2018), rs16985478 (Yu et al. 2018), rs1061684 (Kuroki et al. 2005; Wiśniewski et al. 2015), rs16985478 (Wiśniewski et al. 2015), and rs41548213 (Wiśniewski et al. 2015). Additionally, the majority of the SNVs identified in this study are located at D1-D4 domains. Variations in Ig-domains are the most likely to affect binding to class-I HLAs (Chapman et al. 2000), highlighting the importance of this kind of genetic diversity evaluation.

For *LILRB2*, we identified 41 variable sites (Table 2), 20 (48.8%) of them showing MAF > 1%, and 34 (58.5%) corresponding to missense variants, with 3 variants not following HWE (rs7247025, *p*-value=0.0247; rs7247055, *p*-value=0.0429; rs10404685, *p*-value=0.0267). Among these variants, only three were previously described in the literature: rs383369 (Hirayasu et al. 2008; Bylińska et al. 2018), rs7247538 (Bylińska et al. 2018), and rs386056 (Hirayasu et al. 2008). One of them, the SNP rs383369 (G>A), located at the signal peptide region, has shown the G allele associated with low LILRB2 expression levels (Hirayasu et al. 2008).

*LILRB2* presented 55 haplotypes, 11 of which with frequencies of at least 1% (Table S2). The *LILRB2* network (Figure 3) reveals the presence of four haplogroups that are defined by many missense variants: rs1128646, rs7247451, and rs7247538 (haplogroup a); rs386056 and rs383369 (haplogroup c); rs7247025, rs28405793, rs16027944, and rs7247055 (haplogroup d). Generally speaking, the variants differentiating the haplogroups (a, c, d) are part of the same LD block and are in high or complete LD with each other (r^2^ > 0.5) (Figure 4). There are three more frequent haplotypes (a1, b1, and c1), which summed correspond to 63.32% of the haplotypes identified in our population (Table S2). Interestingly, the SNPs differentiating the haplogroup a (rs1128646, rs7247451, and rs7247538) correspond to positively selected residues according to FUBAR analysis, with rs1128646 and rs7247538 having still been classified as stabilizing mutations by FoldX4. Haplotype c1 is the only one harboring reference alleles for rs386056 (*T) and rs383369 (*G). The rs386056 codifies a residue located at the D2 domain, with the presence of the rs386056*G at this position causing a slight destabilizing effect according to FoldX4. The rs383369*G corresponds to an allele previously associated with LILRB2 expression levels (Hirayasu et al 2008). It should be emphasized that LD is often population-specific, being affected by factors such as non-random mating, selection, mutation, migration or admixture, genetic drift, effective population size, and epistasis (Sabeti et al. 2006). Therefore, the allelic association observed for *LILRB1* and *LILRB2* genes may be influenced by these different processes in a population-specific manner (Pritchard and Przeworski 2001; Flint-Garcia et al. 2003), suggesting that such conclusions may not be directly extrapolated to worldwide populations without a comprehensive evaluation.

*HLA* genes present LD levels higher than expected (Hedrick 1986), which result from natural selection action favoring specific combinations of alleles on the same haplotype (Garrigan and Hedrick 2003). We found reasonable LD differences between both genes, with *LILRB1* presenting four LD blocks well distributed along the gene region, with an apparent disruption of LD between the third and fourth blocks, and with *LILRB2* presenting four LD blocks concentrated in the first half of the gene, between exons 3 and 7, almost without a LD break within the gene. Interestingly, within the *HLA* region, the most significant deviation from neutrality has been observed for class-I haplotypes, which could be due to the requirement for these proteins to recognize a wide repertoire of pathogens and receptors. Likewise LILRB1/2, the distribution of *HLA* alleles and haplotypes may indicate that epistatic effects impact how selection operates on the HLA system, with the epistasis reinforcing LD by selection against recombinants (Alter et al. 2017). Once that epistatic selection exceeds the recombination rate, it can prevent the breakdown of LD and eliminate recombination hotspots. Epistatic selection generally acts against recombinants because they may have a lower fitness due to the sharing of deleterious mutations when compared to both parental haplotypes (Van Oosterhout 2009).

### Misassessment of worldwide *LILRB1* and *LILRB2* variability in the 1000 Genomes data

Data from 1000 Genomes Project provide a broad reference for the comprehension of evolutionary and demographic processes and have been widely used for estimating allele frequencies, phasing, and data imputation. Therefore, we employed the 1000 Genomes data to compare the results found in the Brazilian population to the worldwide population’s data. Considering *LILRB1*, 238 variants were found, with 72 of them corresponding to singletons, removed from analysis. For *LILRB2*, it was found 215 variants, 45 singletons, thus, remaining 170 variants to be evaluated in the subsequent analysis. Additionally, a large proportion of variable sites described in the 1000 Genomes populations did not fit HWE. On average, around 32% of *LILRB1* SNVs and about 14% of *LILRB2* SNVs showed large deviations from HWE expectations, with *p*-values as low as 6.63 x 10^−141^. Some variants displayed heterozygosis unreasonably larger than the maximum expected theoretically. For example, the *LILRB2* variant rs28405793 presented a completely unreliable heterozygosis of 95.8% (*p*-value = 2.03 x 10^−92^) in the European population. A more comprehensive analytical method is necessary before performing additional comparisons involving 1KGen data.

As it has been already demonstrated by Brandt et al. (2015), the mapping approach used by 1000 Genomes Project should be reevaluated in those cases where the studied genes present high sequence similarity with their other family members (e.g., gene families such as *KIR, HLA*, and *LILRB* genes). The mapping procedure involving closely related sequences increases the chance that a read will map to two or more genomic regions. Consequently, such mapping bias results in both the identification of spurious variation sites and in highly biased allele frequency estimations for true variation sites. Therefore, we have decided to disregard the 1000 Genomes data, preventing its use in subsequent comparisons aiming to provide a worldwide perspective on *LILRB1* and *LILRB2* genetic diversity. Moreover, this comparison makes clear the need for a customized method for *LILRB* gene evaluation.

### Evolutionary and structural insights from *LILRB1* and *LILRB2* variability

It is noteworthy that the genetic variability found here is consistent with previous studies suggesting that *LILR* genes are highly polymorphic (Young et al. 2001; Papanikolaou et al. 2004; Kuroki et al. 2005; Hirayasu et al. 2008). Additionally, we observed a high proportion of missense mutations in both genes, which is in agreement with the previous assessment that 58% of the 13,000 exonic SNPs per person are nonsynonymous (Tennessen et al. 2012). Missense variants usually have a high impact on protein activity, affecting processes such as folding, ligand binding, allosteric regulation, post-translational modification, and protein-protein interactions (Stefl et al. 2013; Yates and Sternberg 2013). However, most of the missense SNPs identified so far have no correlation with phenotypic features, compromising their clinical utility. The strategy used in this study, combining population genetics data with protein structure stability analysis, provides insights into the differential impact of *LILRB1/2* wild type or alternative missense variants on the protein structure and function.

### *LILRB1* insights

Except for exons 3, 4, 5, other *LILRB1* segments displayed a significant heterozygosity deficiency compared to neutrality expectations, which is consistent with the directional selection hypothesis (Table 3). Considering Tajima’s *D* test (Table 3), all comparisons showed significant deviations from the neutral equilibrium model. Except for exons 8, 9, 10, which presented a positive *D* value, which is indicative of balancing selection, all other regions presented a negative *D* value, reflecting an excess of rare variants, consistent with either positive or purifying selection. These results point to a predominant effect of either positive or purifying selection over *LILRB1*. Notwithstanding that, the *d*_*N*_/*d*_*S*_ test (Table 4) failed in confirming such conclusion. However, the FUBAR method, designed to detect codon-based signatures of selection, identified nine amino acid residues under positive selection (Table 5). Many of these residues are located at the final portion of the gene, at the cytoplasmic tail. Others are within the Ig-domains: rs1061679, which is known to affect the LILRB1 protein folding (Davidson et al. 2010), rs1061680, and rs1061681, in which the presence of the replacement I132S influences the binding of LILRB1 to HLA-G (Yu et al. 2018). Moreover, FUBAR identified eight residues along the *LILRB1* under purifying selection, all corresponding to synonymous variants. Nonetheless, only nonsynonymous variants were considered for the structural analysis, given that only these mutations can modify the biophysical properties of the encoded protein (Ancien et al. 2018). Among the LILRB1 positively selected residues (Table 5), P68L and T142I, as suggested by FoldX4 (Figure 5), and I155S, as identified by both FoldX and ENCoM (Figure 7), probably affect protein stability; in fact, these positions were previously related to structural stability or protein folding (Davidson et al. 2010; Yu et al. 2018).

Nonsynonymous SNPs located in the LILRB1 D1D2, P68L (rs1061679), T93A (rs12460501), T142I (rs1061680), and I155S (rs1061681) configure a major haplotype that encodes the amino acids LAIS and several others haplotypes that had been selected for *in vitro* assays, showing that this set of mutations have no significant structural changes in receptor conformation; however, it was observed a stronger binding of PTTI with HLA-Cw15 and HLA-B58 when compared to LAIS (Yu et al. 2018). In this context, previous works found no difference when testing these haplotypes, which the authors attribute to the fact that these residues are not in direct contact with HLA-class I or β2m (Kuroki et al. 2005).

It has been proposed that the substitution of proline to leucine at position 68 could interfere with protein structure and conformational dynamics once that proline introduces rigidity into the backbone and leucine is a larger hydrophobic residue (Yu et al. 2018). In this regard, P68L might perturb protein folding (Davidson et al. 2010) and to reduce binding to HLA-G (Yu et al. 2018). This is in agreement with our FoldX4 result that showed that this modification produces a moderate destabilizing effect on the protein.

Located in the D2 domain, T142I has already been evaluated in HCV and HIV context, being associated with both HCMV disease and low CD4 + T cells count (Affandi et al. 2012). It has been shown that 142Isoleucine significantly reduces the interaction with HLA-G, UL18, and Cw15 (Yu et al. 2018), with the presence of threonine at this position being required for glycosylation of the N117 residue, which influences ligand binding (Kuroki et al. 2005). 142Threonine had a slightly stabilizing effect predicted by FoldX4 (Figure 5). Therefore, our analysis agrees with the finding that the presence of threonine at residue 142 induces higher glycosylation, which is recognized as having a significant impact on the stability and protein structure, by enhancing the thermal and kinetic stability of these proteins (Shental-Bechor and Levy 2008).

A previous study has shown that the introduction of a serine at residue 155 significantly increased the interaction of LILRB1 with HLA-G (Yu et al. 2018). FoldX4 has found that I155S is the most strongly destabilizing LILRB1 mutation, with ΔΔG=4.36 (Figure 5). Likewise, ENCoM has shown that I155S produces a destabilizing effect on other D2 residues, leading to higher protein flexibility (Figure 7). Despite appearing contradictory, the greater flexibility when 155Serine is present could lead to a higher interaction with HLA-G since that protein function requires an appropriate balance between molecular stability and structural flexibility (Fields 2001).

Other residues under positive selection (Table 5) have not been identified by this structural analysis because of the lack of an available structure for this purpose, since the remaining positively selected sites are located outside the LILRB1 D1/D2/D3/D4 domains.

The A93T was not identified as positively selected using FUBAR, but the FoldX4 analysis indicated it as the second most destabilizing LILRB1 residue (Figure 5). This residue is located in the region that connects D1 and D2 in LILRB1, having been suggested that A93T may impact the hinge region (Yu et al. 2018).

Although there is no reference in the literature about it, L114R, L197P, and G350R were identified by FoldX4 analysis as strongly destabilizing mutations (Figure 5). Meanwhile, L114R induced a stability increase on other LILRB1 residues, but not on itself, according to ENCoM analysis (Figure 7). Therefore, we hypothesize that a hydrophobicity decrease, according to the Kyte-Doolittle scale (Kyte and Doolittle 1982), might explain the destabilizing effect of these mutations. This hypothesis follows the general idea that hydrophobic effects, i.e., the tendency of non-polar solutes to cluster in water, are the major forces involved in initializing protein folding and stabilizing the three-dimensional structures of proteins (Dill 1990; Pace et al. 2011; Kazlauskas 2018). Once that folded state of a protein is often its lowest energy conformation, the chains are oriented to maximize the hydrophobic effect and make hydrogen bonds and other electrostatic interactions (Kazlauskas 2018).

### *LILRB2* insights

Considering *LILRB2*, the Ewens-Watterson test (Table 3) showed a heterozygous deficiency pattern, consistent with the directional selection hypothesis. Tajima’s *D* test presented negative and statistically significant values for all comparisons made, indicating either positive or purifying selection (Table 3). The *d*_*N*_/*d*_*S*_ test revealed a signal for positive selection at exons 3, 4, 5, 6 (Table 4). FUBAR identified five residues showing evidence of positive selection (Table 6), three of them located at exon 6, exactly where we observed the single significant *d*_*N*_/*d*_*S*_ result. Evidence for positive selection in *LILRB2* has been previously described (Hirayasu et al. 2008). Four residues showing purifying selection signals were identified along the *LILRB2* region, including synonymous variants with no effect in the protein structure.

To the best of our knowledge, no study has evaluated how *LILRB2* polymorphisms influence interaction with HLA molecules. Three of the positively selected LILRB2 residues were also identified as affecting the protein structure by FoldX analysis (Figure 5): T318A, H300Y, and R322P. While T318A was classified as a slightly destabilizing mutation, H300Y and R322P were identified as increasing the protein stability (Figure 5). Except for R322P, in which a change of hydrophilic amino acid for a neutral one occurs, we could not explain these structure alterations based on hydrophobicity effects. Additionally, the mutations with higher effect in structure stabilization (Figure 5) are not under selective pressure according to FUBAR analysis, neither are located in interaction sites with HLA, being located in D3/D4 domains and extracellular portion: I317A, F326S, and P375A/T.

ENCoM identified only R103H as affecting itself (Figure 9). Meanwhile, FoldX4 classified it as a neutral mutation (Figure 5). Two of the other LILRB2 residues identified as being under positive selection, R349W and C306W, were classified as neutral by both FoldX4 and ENCoM.

### Summary findings

Considering both genes, according to the FoldX4 analysis, a higher proportion of mutations affecting the given residue was observed for LILRB1 (65.2%) when compared to LILRB2 (47.6%) (Figure 5). Moreover, it was found that LILRB1 shows lower average stability than LILRB2 (Figure 6). Finally, despite the lack of many residues in the crystalline LILRB1 structure, we found a higher number of LILRB1 residues either under positive selection or affecting the structure protein when compared to LILRB2; however, these residues are not located in interaction sites with HLA-G.

LILRB1 and LILRB2 recognizes HLA molecules (in complex with ß2m) on natural killer (NK) and other immune cells, being these receptors highly polymorphic, with mutations that may affect their activity. In this study, we identified five highly destabilizing mutations in LILRB1 structures using FoldX4, with four of them located in D1 and D2 domains (Figure 5). It is also known that D1 interacts with ß2m and α3 in HLA molecules, and D2 interacts with ß2m (Figure S1). Although LILRB2 also interacts with HLA (Figure S2), there is only one mutation in D1 (R103H), which was considered neutral by FoldX4, and one variant in D2 (M235V) that was classified as slightly destabilizing by FoldX4. Indeed, we identified six other destabilizing mutations, and two stabilizing variations in LILRB2, all of them located in D3 and D4 domains or extracellular portion (Figure 5).

Since *LILRB1/2* modulates the immune response through tight interaction with HLA class-I and other molecules, we may expect a selective regimen favoring the maintenance of a conserved protein structure is expected. Therefore, the purifying selection signal acting over these genes may help eliminate new mutations that interfere with the protein binding through affinity interaction changes. Meanwhile, positive selection may promote the persistence of variations that result in a differential binding ability. The majority of residues identified under positive selection in LILRB1 and all residues identified as such in LILRB2 are located outside the D1/D2domains. For LILRB2, these residues are mainly located at D3/D4 domains. Although it had been shown that D3/D4 domains do not participate in HLA-I binding, they act as a scaffold for mediating binding by D1/D2 (Wang et al. 2019). Therefore, these findings are in agreement with the hypothesis that there would be a higher effect of purifying selection, with sequence conservation in binding regions D1/D2. At the same time, residues in D3/D4 would be more prone to a positive selection effect.

Although most nonsynonymous variations are removed by purifying selection, genes with central functions tend to evolve at higher rates, with some of them showing recent positive selection signals (Eyre-Walker et al. 2006; Harris and Meyer 2006). Considering that *LILRB1/2* are HLA class-I recognizing receptors, these positively selected variations and their effects on the biophysical properties of the encoded proteins might be important for their function, i.e., the differential binding with these ligands, and, therefore, directly influencing pathogen-host interactions.

## Conclusions

This study evaluated, using massively-parallel sequencing, the genetic diversity of *LILRB1/2* coding regions in an admixed Brazilian population sample. Coding *LILRB1/2* regions present reasonable LD levels, as is observed for its main ligand HLA-G. Taken together, as expected, these data provide evidence that natural selection has shaped *LILRB1* and *LILRB2* allele frequencies in the Brazilian population. Furthermore, our structural analysis findings suggest an influence of allelic variation on stability conformational of LILRB1 and LILRB2 molecules. Notwithstanding, further studies are required for determining the combined effect of these mutations in binding among these proteins and its HLA ligands.

## Availability Statement

The genetic data is available upon request.

## Acknowledgements

This work was supported by the Coordenação de Aperfeiçoamento de Pessoal de Nível Superior – Brasil (CAPES) – Finance Code 001; CNPq/Brazil (Conselho Nacional de Desenvolvimento Científico e Tecnológico) (grant number 448242/2014-1); FAPESP/Brazil (Fundação de Amparo à Pesquisa do Estado de São Paulo) (grant numbers 2013/15447-0 and 2017/19223-0); and by the Brazil-France research cooperation program USP/COFECUB (grant no. Uc Me 169-17). C.T.M.J. (312802/2018-8), E.A.D. (304931/2014-1) and E.C.C. (304755/2019-2) are supported by Research fellowships from CNPq/Brazil (Conselho Nacional de Desenvolvimento Científico e Tecnológico).

We thank Juliana Doblas Massaro, Sandra Rodrigues, and Flavia Tremeschin de Almeida for the technical assistance. We also thank Jennifer Thalita Targino dos Santos, Ana Paula Torres, and Marina Tomazela for help collecting blood samples from volunteers.

## Authors’ contributions

MLGO, ECC, SG and CTMJ conceived the study; MLGO, LM, AEP, TMTC, prepared the samples and performed the sequencing experiments, ECC, LCVC, ASS and HSA analyzed the primary data; MLGO, ECC and CTMJ performed the statistical analysis. SG performed the structural analysis. MLGO, ECC, SG and CTMJ interpreted and discussed the results; MLGO and SG wrote the first manuscript draft, which was revised by ECC, ALS, EAD, DC, AS, SG and CTMJ. The final version was approved by all coauthors.

